# Molecular determinants of TRAF6 binding specificity suggest that native interaction partners are not optimized for affinity

**DOI:** 10.1101/2022.05.08.491058

**Authors:** Jackson C. Halpin, Dustin Whitney, Federica Rigoldi, Venkat Sivaraman, Avinoam Singer, Amy E. Keating

## Abstract

TRAF6 is an adapter protein and E3 ubiquitin ligase involved in signaling downstream of cell receptors essential for development and the immune system. TRAF6 participates in many protein-protein interactions, some of which are mediated by a C-terminal MATH domain that recruits TRAF6 to cell-surface receptors and associated proteins. The TRAF6 MATH domain binds to short peptide segments containing the motif PxExx[FYWHDE], where x is any amino acid. Blocking TRAF6 interactions is associated with favorable effects in various disease models. To better define the TRAF6 MATH domain binding preferences, we generated a bacterial cell-surface peptide display library to sample the TRAF6 motif sequence space. We performed sorting experiments and identified 236 of the best TRAF6-interacting peptides and a set of 1,200 peptides that match the sequence PxE but do not bind TRAF6. Selected binders, tested by single-clone bacterial display titrations and bio-layer interferometry, bound TRAF6 tighter than previously measured native peptides. To elucidate the structural basis for TRAF6 interaction preferences, we built all-atom structural models of the TRAF6 MATH domain in complex with high-affinity binders and motif-matching nonbinders that were identified in the screen. We identified motif features that favor binding to TRAF6 as well as negative design elements distributed across the motif that can disfavor or preclude binding. Searching the human proteome for matches to the library screening-defined binding motif revealed that most known, biologically relevant TRAF6 motif matches occupy a different sequence space from the most enriched hits discovered in combinatorial library screening. Our experimentally determined binding preferences and structural models can support the design of peptide-based interaction inhibitors with higher affinities than endogenous TRAF6 ligands.

## INTRODUCTION

Protein-protein interactions assemble signal transduction networks that are critical for cellular function. Knowledge of which proteins interact, and how, is essential for a mechanistic understanding of information propagation through cells. Such an understanding would allow improved network modeling and provide insights into the logic of signal propagation. A high-resolution map of signaling protein connectivity would also advance efforts to identify interactions that can be targeted for therapeutic benefit and provide leads for developing selective inhibitors.

Many interactions important for signaling involve the binding of a recognition domain in one protein by a short linear interaction motif (SLiM) in a partner. Many such domain/motif pairs, including the TRAF6 MATH domain/TRAF6-interaction motif pair that is the subject of this work, have been compiled in the Eukaryotic Linear Motif database [1]. Most motif definitions are based on patterns found in a few examples, leading to incomplete models that do not fully capture the sequence features necessary or sufficient for binding in the cell. A deeper understanding of SLiM sequence requirements can come from large-scale screens, which can provide a more comprehensive view of protein recognition domain specificity [2–7].

TRAF6 is a member of the Tumor Necrosis Factor Receptor Associated Factor (TRAF) family of adaptor proteins with E3 ubiquitin ligase functions [8–10]. TRAF6 mediates NF-κB signaling and thereby participates in immunity and inflammation-related pathways. TRAF6 binds directly or indirectly to Tumor Necrosis Factor receptors and members of the interleukin-1 (IL-1) receptor/Toll-like receptor superfamily, among other proteins. Downstream targets for TRAF6-mediated K63-linked ubiquitylation connect to the regulation of proteins such as transforming growth factor-β-activated kinase-1 (TAK1), IκB kinase (IKK), and mitogen-activated protein (MAP) kinases, which subsequently lead to the regulation of NF-κB and AP-1 activity [9,11]. Direct inhibition of the C-terminal domain of TRAF6 (the TRAF-C Meprin and TRAF Homology – or MATH – domain) has been proposed and explored as a potential therapeutic strategy for the treatment of a variety of pathologies such as cardiovascular diseases, diseases associated with obesity, osteoporosis, and others. [12–18].

TRAF6, like other members of the TRAF family, has four domains. The N-terminal RING domain works in concert with the immediately C-terminal zinc finger domains as an E3 ubiquitin ligase. A coiled-coil domain trimerizes TRAF6. The 17.4 kDa C-terminal MATH domain engages peptides containing TRAF interaction motifs (TIMs) and is responsible for cellular localization [19]. The MATH domains of TRAFs 1, 2, 3, and 5 share high sequence similarity, whereas TRAF4 and TRAF6 are more diverged in sequence and function [9,20–22]. TRAF6 MATH is reported to bind peptides that contain the motif xxxPxExx[FYWHDE] (here referred to as TIM6; Figure 1A), where x is any amino acid [19,23–26]. TRAFs 1, 2, 3, and 5 have been shown to bind [PSAT]x[QE]E, PxQxxD, and PxQxT motifs [1]. For TRAF6, the proline and glutamate residues, referenced here as motif positions (0) and (+2), appear strictly conserved for TRAF6 binding [19,23–26]. A preference for aromatic or acidic residues at (+5) is maintained in the peptide sequences that have been experimentally validated to bind to the TRAF6 MATH domain (Figure 1A) [19,23–26]. Solved structures show that the TRAF6 MATH domain binds to peptides, including the TIM6 motif in CD40, through an interaction that involves the extension of a beta-sheet in the MATH domain (Figure 1B-D) [19,23,24]. Residues in position (+5) bind in a pocket comprised of aromatic and basic residues, engaging in electrostatic and pi-pi interactions (Figure 1D).

**Figure 1.**
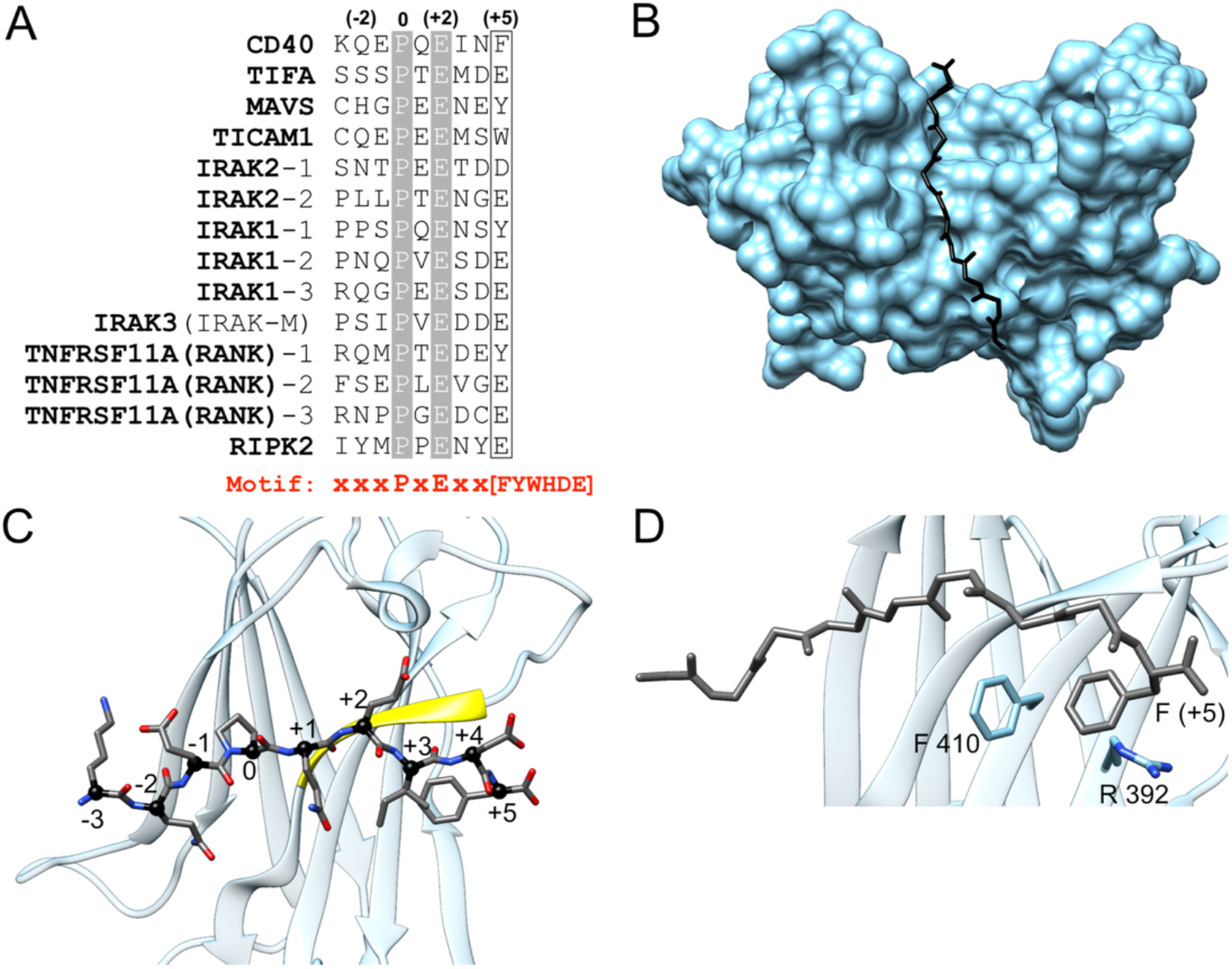
TRAF6 MATH domain interactions with TIM6 peptide ligands. (A) Alignment of TRAF6-binding sequences from known partners showing the numbering scheme used throughout this paper. (B-D) Structure of the TRAF6 MATH domain (cyan) bound to a peptide from CD40 (dark grey), PDB ID 1LB6. (B) MATH domain in surface representation bound to the CD40 peptide. (C) Bound peptide with positions numbered as in (A). Peptide residues at (+1) - (+5) form a beta-strand that pairs with the MATH domain (paired strand in yellow). (D) Interaction of the (+5) Phe in CD40 with Phe 410 and Arg 392 in TRAF6.

Given the low complexity of the TRAF6 motif PxExx[FYWHDE], we reasoned that there might be other determinants of high-affinity TRAF6 binding. To define motif-proximal features important for the biochemical interaction of SLiMs with the TRAF6 MATH domain, we used bacterial surface-display screening to explore sequence preferences within a combinatorial library denoted xxxPxExxx, with x being a random amino acid, keeping the proline fixed at position (+0) and the glutamate fixed at position (+2). We screened this library and identified 236 highly enriched binders and 1200 nonbinders. We then used structure-based modeling to explore the nature of interactions between the best-binding peptides and the MATH domain.

Our analysis revealed residues within the motif that support high-affinity binding and illuminated negative-design elements that explain why many peptides that contain PxE are not suitable TRAF6 ligands. These insights help to elucidate the determinants of TRAF6 binding specificity. We compared the sequence features of the library-identified binders with reported TRAF6 binders and found that most native interaction partners do not match the top sequences isolated from the library. Notably, there are no sequences in the human proteome that share all of the features that are prominent among the tightest binders from the screen. These results suggest that native TRAF6 interaction partners may be under functional selection for moderate affinity, providing an opportunity to out-compete native interactions with designed peptides or mini-proteins.

## RESULTS

### Library screening by cell-surface display reveals strong positional preferences for peptides that bind TRAF6

Bacterial-surface display can provide information about the binding of short peptides to protein interaction domains [3,27]. For this work, we developed bacterial-surface display constructs in which TIM6 peptides were fused to the C-terminus of reengineered OmpX [28,29], such that when the construct was expressed, the TIM6 peptides were presented on the outer membrane of *Escherichia coli* cells. Peptide-displaying cells were incubated with biotinylated TRAF6, and binding was detected using streptavidin conjugated to phycoerythrin by fluorescence-activated cell sorting (FACS). The level of peptide expression was quantified using a FLAG-binding antibody conjugated to allophycocyanin (details in Methods). By titrating TRAF6 MATH domain into a clonal population of peptide-displaying cells, an apparent cell-surface dissociation constant (K_d_) can be determined by fitting the binding signal vs. TRAF6 concentration. We measured peptide binding to TRAF6 homotrimers consisting of the coiled-coil and MATH domains (here the construct is termed T6cc). Binding between TRAF6 trimers and peptide-displaying cells was multivalent, and the associated avidity enhanced the apparent dissociation constants. For example, by ITC, the isolated TRAF6 MATH domain binds to a peptide from CD40 (KQEPQEIDF) with a K_d_ of 83 µM [19]. In contrast, this peptide embedded in our display construct bound to our trimeric TRAF6 construct with an apparent K_d_ of 1.2 µM by bacterial surface display.

To evaluate the TRAF6 MATH domain interaction motif space, we constructed a combinatorial library by introducing random residue variation around the core TIM6 element PxE. We used degenerate NNK codons to encode any of 20 amino acids at “x” positions in the sequence xxxPxExxx. The proline at position (+0) and the glutamate at position (+2) were held fixed to increase the proportion of binders in the library and to force a specific binding register to facilitate analysis and modeling. These sequences were presented in the context of flanking sequences from CD40; see Table S1 for details of the display constructs. To isolate cells displaying peptides that bound to the TRAF6 trimer, we carried out one round of enrichment using magnetic microbeads (see Methods). This procedure generated a smaller library, enriched in TRAF6 binders, that we designate MACSLib; this library was used as the input for subsequent enrichment experiments. Deep sequencing analysis of all unique sequences over all rounds of enrichment revealed that MACSLib contained a minimum of 8000 unique peptides matching the sequence xxxPxExxx. The content of the library is summarized in Figure 2.

**Figure 2.**
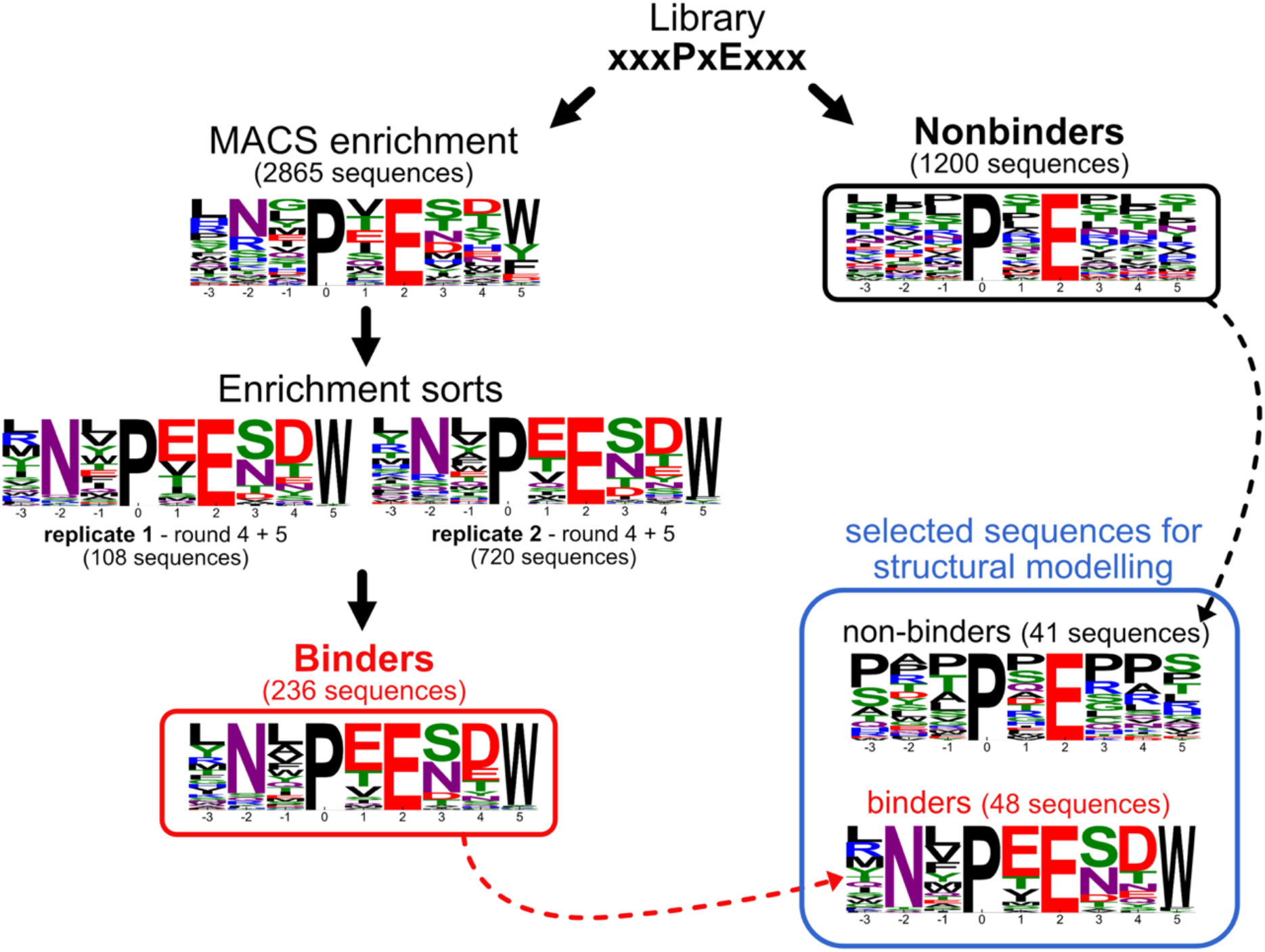
Sequence logos for TRAF6-binding and nonbinding peptides. TRAF6 binders were identified by initial MACS enrichment followed by 2 replicate 5-round FACS enrichment experiments. The final list of binders (red box) was generated by combining both replicates and further filtering for sequences that enriched across multiple rounds. Nonbinder sequences were defined as significantly populated sequences (read count >= 20) in the nonbinder pool (black box). Sequences selected for structural modeling are shown in the blue box. Residue height in the logos represents the frequency of that residue in the sequence set. The number of sequences in each set is shown in parentheses.

To identify high-affinity binders in MACSLib, two separate 5-round enrichment sorts were performed using FACS to separate binding library members from nonbinding members (details in Methods). The stringency of the binding assay was gradually increased by using a lower concentration of T6cc for each round (concentrations used: 300 nM, 100 nM, 30 nM, 10 nM, 3 nM). Following sorting, the population of binding cells in each round was deep sequenced to monitor the enrichment of individual sequences. The two replicate sorting experiments gave similar results, with sequences from rounds four and five reflecting similar preferred residues, indicating convergence of the selection process (Figure 2). In both replicate experiments, the three sequences LNLPEESDW, RNVPEESDW, and TNWPEENDW ranked among the top four binders based on sequencing read counts, and 14 of the top 20 most-represented sequences were the same in the two datasets. Examination of enriched sequences, particularly those in the final rounds, indicated a strong preference for Trp at position (+5). Additionally, preferences were evident for Asn at (−2), Glu at (+1), Ser/Asn at (+3), and a polar or acidic residue at (+4). A final set of 236 high-confidence TRAF6 binders was generated by taking the set of sequences present in rounds 4 and 5 of either replicate and filtering for sequences that enriched over at least 2 rounds of sorting (Figure 2, red box and supplementary data files; see Methods). We also generated a population of nonbinders by collecting cells from the original unenriched library that gave a strong peptide-expression signal but no TRAF6 binding signal. This logo did not show strong enrichment of any particular features (Figure 2, black box).

To verify that the screening hits bound to TRAF6 in a concentration-dependent manner, we performed single-clone titration experiments to measure apparent cell-surface dissociation constants, 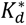 values, for 14 sequences selected from the enrichment data (Figure 6A and Figure S1; see Methods). We also measured 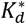 for the native 9-residue CD40 TIM6 peptide in the same context. Interestingly, all of the top peptides from the enrichment bound TRAF6 with an apparent affinity higher than the CD40 TIM6 peptide, with some binding over an order of magnitude tighter than the CD40 TIM6. For example, RNVPEESDW, LNLPEESDW, and TNWPEENDW bound TRAF6 with 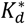 values of 31 nM, 46 nM, and 84 nM, respectively, whereas the CD40 TIM6 bound with 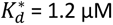. To validate the cell-surface interactions identified in the screen, we measured TRAF6-peptide binding by biolayer interferometry (BLI), using purified monomeric TRAF6 MATH domain (construct termed T6m) in solution and purified peptides attached to the sensor tip (see Table S1 for construct details). By BLI, RNVPEESDW, LNLPEESDW, and TNWPEENDW bound to TRAF6 with solution K_d_ values of 24.0, 27.5, and 37.2 µM, respectively, while the CD40 TIM6 bound with a K_d_ of 238 µM. The BLI data validate the cell-surface display results and support the conclusion that top hits from the screen bind with higher affinity than the TIM6 motif in CD40, which is one of the tightest known TRAF6 peptide binders [19] (Figure S2). Based on these observations, we conclude that despite the screening assay being performed in the environment of the cell surface, and in a multi-valent context, enrichment sorting returned high-affinity binders.

### Structural modeling explains positive and negative binding determinants

For computational analysis of the structural determinants of TRAF6-TIM6 binding, we chose a subset of high-affinity binders identified from enrichment sorting and a subset of sequences designated as nonbinders (see Methods). Figure 2 shows sequence logos summarizing features of these two subsets of sequences. We then tested whether structure-based models could discriminate the binders from the nonbinders.

FlexPepBind (FPB) is a peptide modeling protocol in the Rosetta suite [30]. As input to FPB, we prepared models using the structure of TRAF6 bound to peptide KQEPQEIDF from CD40 (PDB ID 1LB6) as a template [19]. We assumed that all peptides bound in the canonical TRAF6 binding groove with an alignment of PxE to the corresponding residues in peptide KQE**P**Q**E**IDF. Starting from this initial docking position, the binding pose of the peptide was sampled, subject to constraints on the distance between the peptide and the MATH domain, and the lowest interface score over all sampled poses was assigned to each peptide complex (see Methods). Figure 3A shows the ranking of the 48 binders and 41 nonbinders by FPB score, which achieves a good separation of the two populations, with only 5 out of 89 complexes misclassified when using an optimal score cutoff. Native CD40 TIM6 peptide (KQEPQEIDF), scored with the same protocol, gave an FPD interface score of -34.2, which is in the weaker end of the range of binding peptides, consistent with the affinity measurements discussed above.

**Figure 3.**
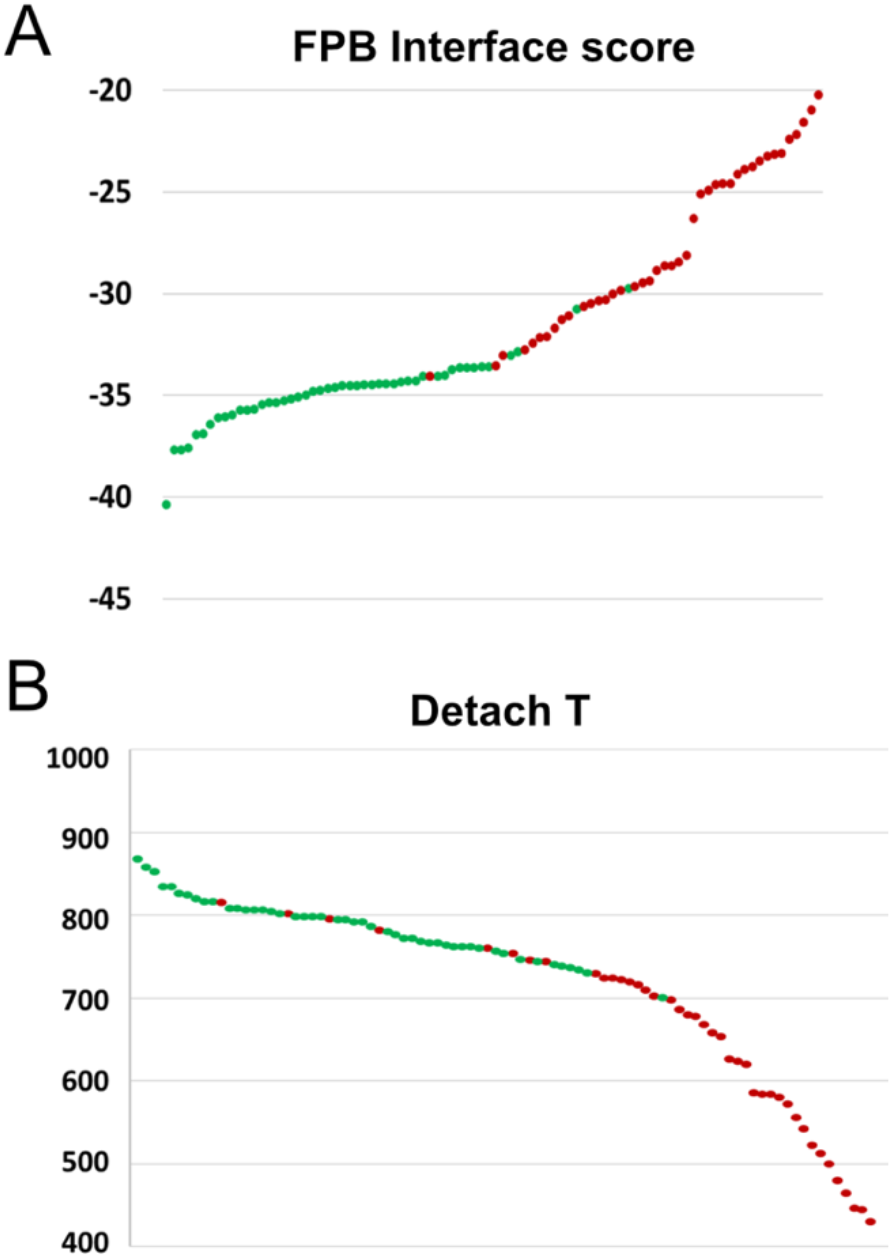
FBD and Detach T scoring of TRAF6 binders and nonbinders. TRAF6 peptide binders (green) and nonbinders (red) identified by high-throughput screening are plotted based on a computationally predicted score: FBD interface score (A) or Detach T (B).

Because 9-residue peptides will sample an ensemble of conformations when bound to the TRAF6 domain, we also tested a molecular dynamics-based protocol for evaluating peptide-domain interactions. Starting with complexes modeled on the structure of TRAF6 bound to KQEPQEIDF, as described above, we computed for each model a detachment temperature (Detach T), corresponding to the temperature at which the distance between the alpha-carbon of TRAF6 Phe 471 and the center of mass of the peptide increased beyond 7 Å, when the temperature was gradually increased from 300 K. Detach T, like FPB interface score, was able to separate binding peptides from nonbinders, as shown in Figure 3B, with no binders giving Detach T values lower than 700 K.

To explore the structural origins of sequence trends apparent from our enrichment sorting results, we used molecular dynamics simulations to analyze TRAF6 complexes with peptide ligands from CD40, RANK, and each of the 48 binders (see Methods). In native complexes and all high-affinity binders, simulations showed the persistence of 5 hydrogen bonds that position the peptide as an extension of the beta-sheet in the MATH domain, as seen for the CD40 peptide in PDB structure 1LB6 (Figures 1C and 4A). The hydrogen bonds involve backbone atoms of residues in positions (+1), (+3), and (+5). These non-covalent interactions were highly stable during all 80 ns of equilibrated-MD simulation for the native CD40 and RANK peptides and for all high-affinity binders. Invariant TIM6 residues Pro at (+0) and Glu at (+2) also preserved their crystallographic positions throughout all trajectories, with only minor displacements (Figure 4B). Pro at (+0) is accommodated in the pocket created by residues Phe 471, Met 450, and Tyr 473, while the negatively charged Glu at (+2) caps a 3-10 helix formed by residues Leu 456, Leu 457, and Ala 458 (Figure 4B) in the MATH domain.

**Figure 4.**
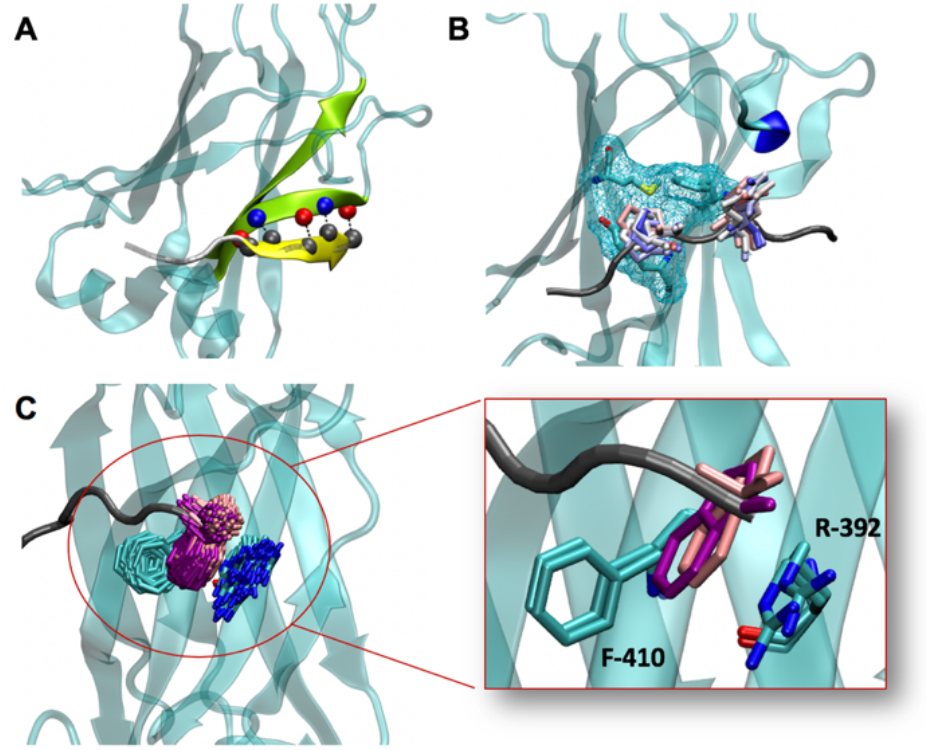
Structurally conserved features of high-affinity complexes. (A) Five beta-sheet hydrogen bonds involve main-chain atoms of residues at positions (+1), (+3), (+5) (yellow), and residues 472, 470, and 468 in the MATH domain (green). The image shows a snapshot from a simulation of TRAF6 MATH in complex with the peptide LNLPEESDW. (B) Positions of Pro at (+0) and Glu at (+2) (sidechains in sticks) from different frames of the equilibrated MD simulation of the peptide LNLPEESDW bound to TRAF6 MATH (color scale: red-white-blue for snapshots from the beginning-middle-end of the equilibrated part of the simulation). Pro binds into the pocket shown with cyan mesh, and Glu caps the short helix marked in blue. (C) The two most populated clusters for Trp conformations at position (+5) for 65% of the high-affinity binders. This sidechain arrangement allows simultaneous pi-pi interaction with Phe 410 and cation-pi interaction with Arg 392. The expanded region highlights two snapshots from the two most common conformations.

Trp at (+5) was present in most of the binders obtained from enrichment sorting, even though this residue is not common in known native interaction partners of TRAF6 (Figure 1A). Indeed, only 14 out of 236 binder sequences identified in the enrichment screen did not have a W at position (+5). Our simulations showed different conformations for the Trp sidechain. Most frequently, the indole group was inserted into the receptor pocket (Figure 4C), allowing for simultaneous pi-pi interaction with Phe 410 and cation-pi interaction with Arg 392. This conformation represented the most populated cluster for 65% of the high-affinity peptides and resembles the conformation for Phe at position (+5) in the complex of CD40 TIM6 peptide bound to TRAF6 MATH (Figure 1D) [19]. In particular, clustering Trp at (+5) conformations from simulation frames by RMSD shows that more than 60% of the conformations are within 1 Å of the sidechain arrangement shown in Figure 4C. We also observed structures in which the Trp indole was flipped out of the pocket but maintained a binding interface, including backbone H-bond interactions with Pro 468 and occasional pi-pi or cation-pi interactions with Phe 410 or Arg 392. Such conformations are shared among 20% of the high-affinity binders. The remaining 15% of high-affinity binders showed unclustered Trp (+5) conformations in which the backbone was still involved in an H-bond interaction with Pro 468, but the indole group was flipped out and did not contact the MATH domain.

The preference for Asn at (−2) in binders from the screen can be explained by its sidechain interaction with nearby Glu 448 on TRAF6 through a hydrogen bond that stabilizes the N-terminal end of the peptide in the pocket (Figure 5A). This interaction is shared by more than 70% of the high-affinity binders, which each form this contact for > 30% of the simulation time. Asn also makes a stable interaction with the backbone of Thr 475 (for > 40% of simulation time for all of the high-affinity binders), which is also apparent in the structure of CD40 TIM6 bound to TRAF6 [19]. Longer residues at (-2) (e.g., Gln) are unable to interact with both TRAF6 amino acids. This interaction pattern appears to be important for high-affinity binding: Asn at (−2) is present in 190 of the 236 binders from the enrichment.

**Figure 5.**
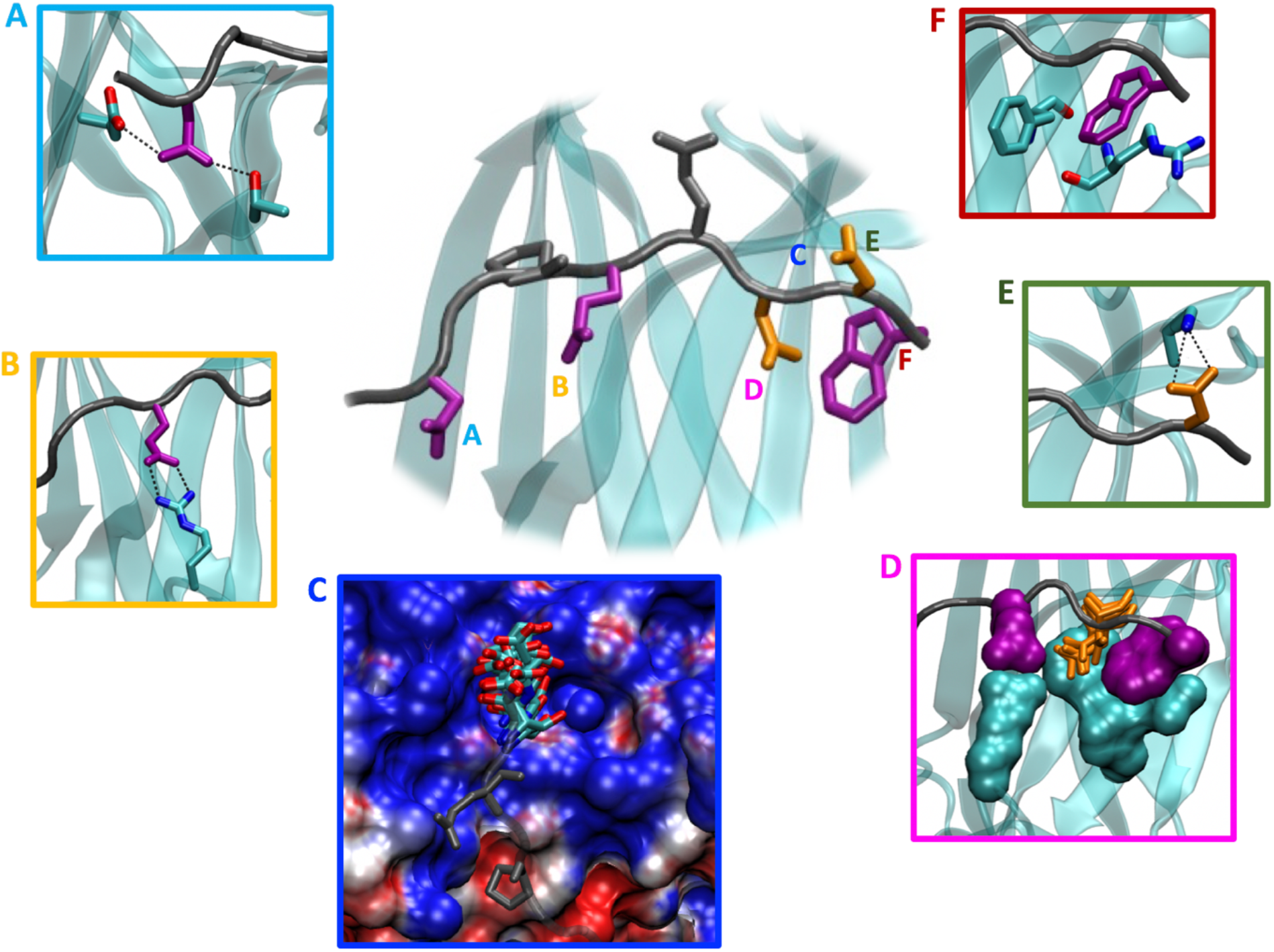
Overview of the most significant contacts between the highest affinity binders and the TRAF6 MATH domain, as captured in molecular dynamics simulations. Panels A – F illustrate specific interactions discussed in the text. Each panel is outlined in a color that matches a label in the central figure. Key peptide residues are represented in sticks: grey for Pro at (+0) and Glu at (+2), purple for residues at positions that favor a particular amino acid, and orange for residues at positions that favor a group of amino acids with similar features. All of the highlighted interactions are present for > 30% of simulation time for all high-affinity binders. (A) Asn at (-2) can simultaneously form H-bonds with Glu 448 and Thr 475. (B) Glu at (+1) forms a bi-dentate salt-bridge interaction with Arg 402 when both sidechains are fully extended. (C) Residues at positions (+3) and (+4), shown here with Asn and Asp in sticks, are located in an electrostatically positive region (as indicated by blue coloring). (D) Interactions involving residues at (+1) and (+5), shown as space-filling Glu and Trp in purple, along with MATH domain residues 402, 410, 392, and 394 (space-filling, cyan), narrow the pocket at position (+3) so that small residues, such as the Asn pictured in orange sticks, are preferred at this site. (E) Residues at position (+4) face solvent, and acidic residues at this site, such as the pictured Asp, can form a salt bridge with Lys 469. (F) Trp at (+5) engages in edge-to-face pi-pi and cation-pi interactions with residues Phe 410 and Arg 392, respectively.

A preference for Glu at position (+1) can be explained by the salt-bridge interaction that this residue makes with Arg 402 in the MATH domain (Figure 5B). Despite a Cα-Cα distance of ∼10 Å, the Arg 402 sidechain can make a salt bridge with the (+1) Glu when both are fully extended toward one another. This interaction is stable for more than 80% of equilibrated trajectory time and is completely missing when Glu is substituted with Asp due to the shorter sidechain of the smaller residue.

At position (+3), interactions involving residues at (+1) and (+5), and the positions of MATH domain residues 392, 394, 402, 410, and 474, narrow the pocket so that small residues, such as Asn, are preferred at this site (Figure 5D). Residues at position (+4) are less sterically constrained and can form a hydrogen bond or salt bridge with surface Lys 469 (Figure 5E); all high-affinity peptides with Glu at (+4) form a salt bridge with Lys 469 in more than 60% of simulation frames. Overall, the surface of TRAF6 is electrostatically positive near positions (+1) to (+5), as shown in Figure 5C.

Analyzing our models of the high-affinity peptides helped explain why many nonbinders did not form tight interactions with TRAF6, despite including the conserved Pro at (+0) and Glu at (+2). At positions (+1), (+3), and (+5), high-affinity binders make hydrogen bonds that complete a beta-sheet with TRAF6. Proline residues are disfavored in beta structures because they lack the required NH group for this interaction and prefer backbone dihedral angles far from the typical range in β-sheets [31]. Thus, Pro at any position between (+1) and (+5) is expected to be highly unfavorable. 386 of the 1200 nonbinders identified in the screen have such a substitution, which is likely sufficient to prevent high-affinity binding. None of the 236 binders contain a proline at these positions. Furthermore, the TRAF6 MATH domain is electrostatically positive near positions (+3), (+4), and (+5) (Figure 5C), suggesting that positively charged residues would be destabilizing at these sites. Indeed, Arg, Lys, or His are found at one or more of these positions in 436 of 1200 nonbinders but in only 6 of 236 binders. Steric constraints at position (+3) are further expected to disfavor medium or large residues at this site. Consistent with this, residues Q, H, I, L, F, Y, or W are found at position (+3) in 396 of the 1200 nonbinders but in only 4 of the 236 binders. Overall, 898 of the 1200 nonbinders have at least 1 of the unfavorable sequence features described above (see Table 1 for summary). The nonbinders also lack key residues that form stabilizing interactions in the highest affinity binders. Only 119 of the 1200 nonbinders contain Asn at (-2), Glu at (+1), or Trp at (+5), while 234 of 236 binders contain at least one of these interactions. Only 3 of 1200 nonbinders contain two or more of these stabilizing residues, while 217 of 236 binders contain two or more of these stabilizing residues.

**Table 1.**
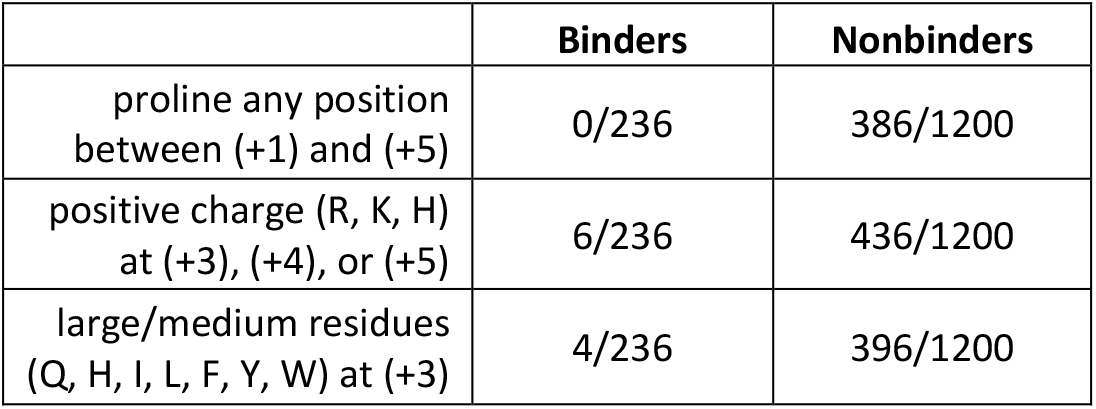
Fraction of binder and nonbinder sequence sets with given sequence features

|  | **Binders** | **Nonbinders** |
| --- | --- | --- |
| proline any position between (+1) and (+5) | 0/236 | 386/1200 |
| positive charge (R, K, H) at (+3), (+4), or (+5) | 6/236 | 436/1200 |
| large/medium residues (Q, H, I, L, F, Y, W) at (+3) | 4/236 | 396/1200 |

### Candidate TRAF6 interaction motifs in the proteome do not share the sequence features of the top screening hits

We investigated whether any human proteins contain close matches to the high-affinity sequences identified by screening. We defined a position-specific scoring matrix (PSSM) to score candidate TRAF6 interaction motifs based on how well they match our top binders. We used pLogo [32], a log-odds-based method, to construct the PSSM, using the 236 binding sequences from the enrichment as the foreground and the 1200 nonbinder sequences as the background. The nonbinder sequences were considered a fair approximation of the sequence composition of the input library, assuming that TRAF6 binders are rare in the library. Indeed, we don’t observe any apparent residue preferences in the nonbinder set (Figure 2). To test if the PSSM score of a sequence represents how well that peptide binds to TRAF6 on the cell surface, we scored the sequences used in single clone titrations with our PSSM. Scores were normalized to span from 0 to 1, with 0 being the lowest possible PSSM score and 1 being the highest possible PSSM score. We found that PSSM score is correlated with apparent cell-surface affinity, suggesting that our model is a good predictor of TRAF6 binding (Figures 6A and B).

**Figure 6.**
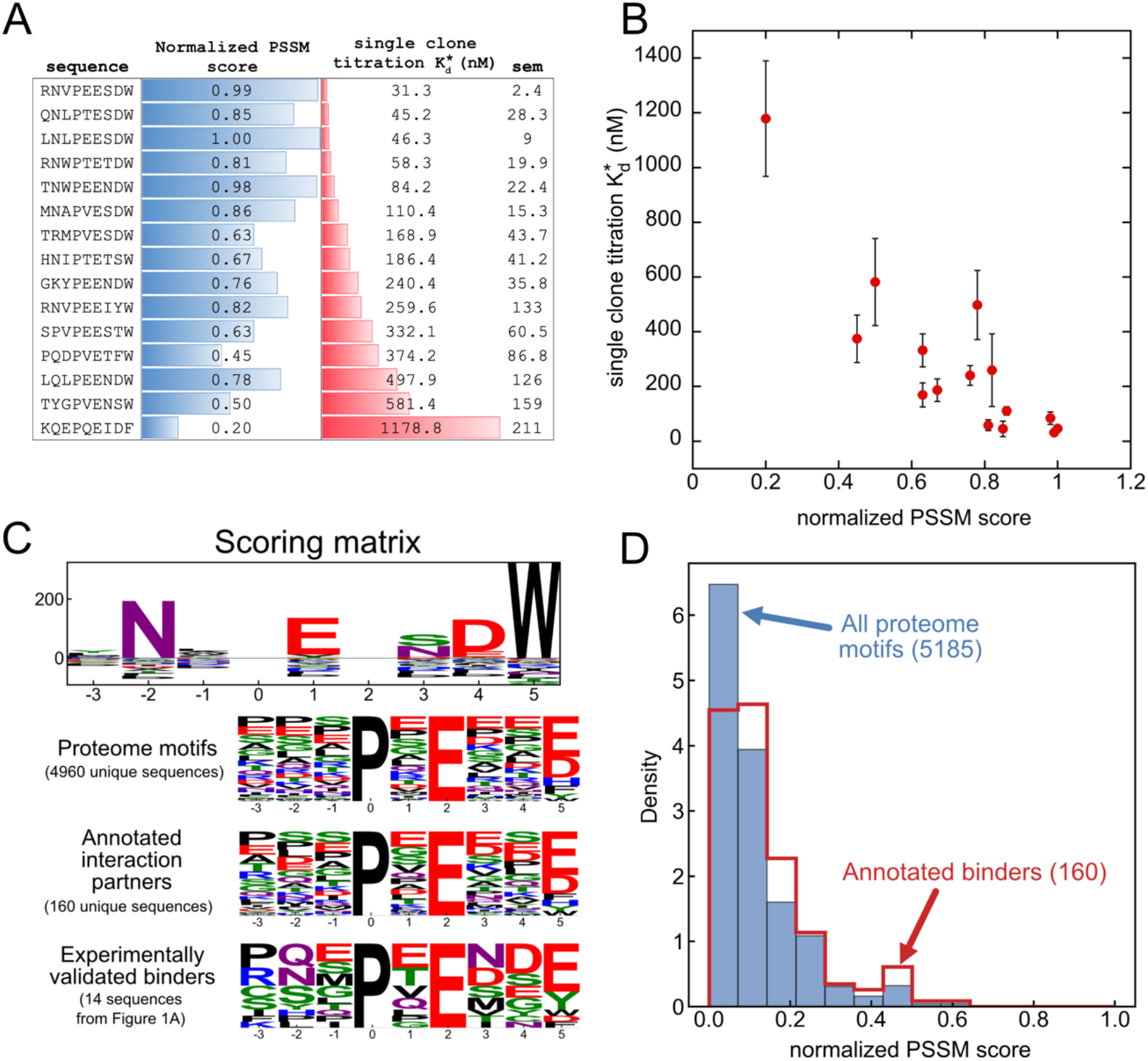
TRAF6 motif scoring. (A and B) The PSSM scores of selected TRAF6 binding peptides are roughly correlated with their apparent cell-surface binding affinity for TRAF6. (A) Reported dissociation constants are the average of fits to 2-3 replicate titrations. The standard error of the mean (sem) is reported for each sequence. (B) Correlation between 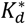and PSSM score (data from A). Error bars are the standard error of the mean of 2-3 replicates. (C, top) Position-specific scoring matrix generated from the screening data and used to score candidate binding motifs in panel D. (C, bottom) Sequence logos of all unique TRAF6 motif matches (motif: xxxPxExx[FYWHDE]) in disordered regions of the proteome (IUPred score > 0.4), compared to the subset of those motif matches within proteins that are annotated as TRAF6 interaction partners (HIPPIE database [35]) as well as experimentally validated TRAF6 MATH domain binders. (D) Distribution of normalized PSSM scores of TRAF6 motif matches in disordered regions of the proteome (IUPred score > 0.4). Scores for all motifs are shown in blue, and motifs within proteins that are annotated as TRAF6 interaction partners (HIPPIE database [35]) are shown in red.

The PSSM was used to score TRAF6 motif matches in the human proteome to identify those SLiMs most likely to bind with high affinity. To focus on regions of the proteome that are predicted to be disordered, the TRAF6 motif matches with an average IUPred score > 0.4 (5185 hits total) were obtained using the SLiMSearch tool [33,34] (regular expression: …P.E..[FYWHDE]). The logo of hits is shown in Figure 6C, along with a logo of the subset of hits that are within proteins annotated to interact with TRAF6 [35], and a logo of experimentally validated TRAF6 binding peptides.

Figure 6D shows the distribution of PSSM scores for the sequences retrieved using SLiMSearch [33]. Notably, no sequences in the proteome occupy the sequence space favored in the screen (i.e., no sequences have a high score), and there is no match to the sequence predicted to be the best TRAF6 binder, YNLPEESDW, anywhere in the proteome. The highest scoring motif with an IUPred score > 0.4 is PNNPQEADW, which has a score of 0.78. In the absence of an IUPred filter, the highest-scoring sequence in the proteome is FNEPEENFW, with a score of 0.85.

Despite the absence of sequences in the human proteome that closely match the sequence space identified in the screen, we nevertheless used the PSSM as one tool for identifying human proteins with characteristics conducive to TRAF6 binding. We constructed a table from the proteome motif matches that includes a variety of scores and filters. Filtering using these scores can restrict candidate motifs in the human proteome to those more likely to interact with TRAF6 (Table S3).

In addition to having good potential for biophysical interaction with the TRAF6 MATH domain, which we assessed using our PSSM, TRAF6-binding SLiMs must be accessible for binding. The hits in our table can be filtered by IUPred score, and we set this to only include hits that are predicted to be disordered (IUPred score > 0.4). However, IUPred score is not a guarantee of accessibility. The AlphaFold pLDDT score is reported to be a good predictor of disorder [36], so we included the average and maximum AlphaFold pLDDT scores of the motif (+3 flanking residues) within the predicted structure of the protein [36–38]. For the average score, we recommend a cutoff of < 65 but caution that this will likely remove some instances where a motif is still accessible, despite the high pLDDT score. The maximum pLDDT score of any residue within the motif +/-3 residues on each side is also reported, as this may detect cases in which most of the motif is disordered but proximity to a folded domain structure limits accessibility. For this filter, we recommend setting it to consider hits with a maximum pLDDT score of less than 70.

We judged that proteins involved in similar biological processes as TRAF6 are more promising candidate interaction partners [33]. To identify MATH domain binding motifs in proteins that share functions in common with TRAF6, we used Gene Ontology (GO) annotations [39,40]. Specifically, we used SLiMSearch to retrieve GO terms for TRAF6, where each term has an associated p-value representing the likelihood that any 2 proteins in the proteome share that term by chance (p-value from SLiMSearch [33]). The table of proteome hits can then be filtered to include proteins that share 1 or more TRAF6 GO terms with a p-value below a given threshold. Smaller p-value cutoffs result in more general GO terms being removed from the list of terms used in the filter and provide greater stringency. We provide several different p-values as options and suggest p=0.01 as a starting point.

Many proteins have been reported to interact with TRAF6 without identification of the mode of interaction. We used the HIPPIE database [35] to identify which of the proteome windows that we evaluated are in proteins that are already annotated to be TRAF6 interaction partners (logo – Figure 6C; score distribution – Figure 6D, red line). A high-scoring motif match in a protein annotated as a TRAF6 interaction partner could indicate that its interaction with TRAF6 is likely to occur through the MATH domain.

Our structural analysis identified several sequence features that disfavor or prevent PxE peptides from binding to the MATH domain (Table 1). We added filters to the table to identify candidate motifs having unfavorable residues, such as a proline at positions (+1) to (+5), a large/medium residue (QHILFYW) at position (+3), or a positively charged residue (RKH) at positions (+3), (+4), or (+5).

Using our table and filters, we identified several interesting potential TRAF6 interaction motifs in the human proteome. One such hit is the sequence GMGPVEESW, which starts at position 350 in RIPK1. The sequence has a Val at (+1), which has a moderately favorable PSSM score, and a highly favorable Trp at (+5). RIPK1 is a serine-threonine kinase involved in regulating TNF-mediated apoptosis, necroptosis, and inflammatory pathways [41]. It has been annotated as a TRAF6 interaction partner in the HIPPIE database, but the details of the interaction are unknown. RIPK1 has been found to bind to other TRAF proteins [42] and also to TICAM1 [43]. We propose that RIPK1 may interact with TRAF6 via the MATH domain, engaging this short segment.

One of the highest-scoring hits in the proteome is the sequence QNFPVESDW (PSSM score = 0.85) from the E3 ubiquitin-protein ligase RNF103. This sequence has the highly favorable residue Asn and Trp at positions (-2) and (+5), respectively. The sequence also contains favorable Ser and Asp residues at (+3) and (+4), respectively, and Val at (+1). RNF103 acts as an E3 ubiquitin-protein ligase that is localized to the ER membrane; it is involved in the ER-associated degradation (ERAD) pathway [44]. The TRAF6 motif match has an average IUPred score of only 0.19 and thus is not predicted to be disordered by this metric. However, the average AlphaFold pLDDT score of the motif is 38.7 (corresponding to predicted disordered) and the residues appear accessible in the AlphaFold-predicted structure [36–38]. Although RNF103 localizes to the ER membrane, the candidate motif (positions 474-482) maps to the cytosol, given its location between the last transmembrane helix and the cytosolic RING domain of RNF103.

Another hit with potential for biological significance is the sequence GNFPEENND, which spans positions 1065 – 1073 in the leptin receptor. This sequence contains an Asn at (-2), Glu at (+1), and Asn at (+3), which are all favorable residues according to our model. The leptin receptor binds leptin, which is secreted from adipose cells. In obese mammals, leptin levels are elevated, leading to chronic low-grade inflammation [45]. TRAF6 is a well-known regulator of the inflammation response, suggesting a link between the two pathways. Follow-up biochemical work accompanied by tests in appropriate cell lines will be required to validate this candidate interaction and the other hits described above.

## DISCUSSION

The discovery of TRAF6 interaction partners over decades of experimental research has led to a general definition of the TRAF6 MATH domain binding motif (xxxPxExx[FYWHDE]). This was arrived at by compiling aligned, verified TRAF6 binding sequences and identifying their common sequence features [19,23–26]. In this work, we explored the TRAF6 motif sequence space more systematically, using cell-surface screening of a combinatorial library that presented the core PxE motif flanked by random residues. The top hits obtained from this screen bound with affinities comparable to or higher than known TRAF6 interaction partners reported in the literature [19,24]. Analysis of screening hits highlighted which residues were most preferred at each position and identified features that differentiate binding sequences from nonbinding sequences among protein segments that contain the core element PxE.

Two different methods of structure-based modeling demonstrated the ability to distinguish the best-binding peptides from the background, and we used molecular dynamics simulations to study the bound ensembles of diverse binders. This analysis provided a structural explanation for the residue preferences observed in our screening data. In particular, in all high-confidence binders that we analyzed, Asn at position (-2) can form favorable interactions with Glu 448 on TRAF6, Glu at position (+1) can form a salt bridge with Arg 402 on TRAF6, and Trp at position (+5) can form pi-pi interactions with Phe 410 and cation-pi interactions with Arg 392. Our structural analysis also highlighted negative design elements that prevent myriad PxE sequences from binding to TRAF6 MATH. The logo of nonbinders in Figure 2 does not indicate any strong features, but our data support a model in which a variety of sequence features, including proline in positions (+1) to (+5), a large residue at position (+3), or a positively charged residue at (+3), (+4), or (+ 5) can disfavor binding. Thus, it appears that within this 7-residue stretch that includes PxE, several positive features and the absence of a variety of negative-design elements are important for making a functional TRAF6 binder.

Our analysis of disordered regions in the proteome revealed that no sequences map to the sequence space that we identified for high-affinity binding TRAF6 motifs using cell-surface display. Indeed, all of the well-studied peptides from verified TRAF6 MATH domain binders lack the features of the highest-affinity binders identified in the screen or only contain 1 or 2 of the most favorable interactions we identified (Figure 1A). Only 3/14 have an Asn at (-2), and only TICAM1 contains Trp at (+5). Our findings imply that the core binding motif is either not under selection for high affinity, or high affinity is detrimental to TRAF6 function.

It has been proposed that peptide recognition domains bind to SLiMs with weak affinity to allow for transient and short-lived interactions [46]. A complex that uses multiple weak interactions rather than one higher-affinity binding site provides opportunities for regulation, as is the case for tandem recognition of SLiMs by SH2, SH3, WW, and other domains [47–50]. The TRAF proteins provide another example of the benefits of weak binding for signaling. TRAF6 uses avidity to significantly enhance binding affinity to oligomeric receptor proteins. Thus, in concentration regimes where the binding affinity of a single-motif peptide is not significant, receptor oligomerization can trigger TRAF6 binding to three or more motifs in the clustered cytoplasmic receptors’ tails, which then promotes ubiquitylation that propagates the signal further downstream [9]. Artificial oligomerization of TRAF6 alone is sufficient to activate signaling through certain pathways [51]. In this scheme, conserving a weak, fast-exchanging interaction between individual motifs and monomeric MATH domains is likely important for supporting rapid, ligand binding-dependent assembly and disassembly of a TRAF6 signaling complex.

When viruses or parasites infect cells, they interfere with host cell regulatory pathways to make the host susceptible to viral replication and control immune response. Many viral/parasite proteins contain motifs that mimic host SLiMs [52]. The requirement to out-compete native SLiM interactions means that viral proteins are often under selective pressure to maintain high-affinity interactions with their host SLiM-binding domains. TRAF6 plays an integral role in immune response pathways and is thus a potential target for viral manipulation. Indeed, several pathogen proteins recruit TRAF6 MATH to modulate the host cell NF-κB pathway, including the protein UL37 from human herpesvirus (motif: SNTPVEDDE) [53] and GRA15 from the intracellular parasite *Toxoplasma gondii* (motif: PQVPGENSY) [54]. Of particular note, the motif in UL37 has residues N, V, and D at (-2), (+1), and (+4), respectively, which are predicted to be favorable for binding to TRAF6 in our model. We suggest that these residues allow UL37 to out-compete native interactions.

High-affinity TRAF6 binders isolated in this work can serve as lead peptides for inhibitor development. TRAF6 signaling is implicated in inflammation and cardiovascular disease [12,13,18]. Targeting TRAF6 MATH is reported to improve insulin sensitivity in obese mice, improve heart function in mouse models of non-ischemic cardiac failure, reduce atherosclerosis, and inhibit osteoclastogenesis and bone resorption [14–17]. A RANK peptide attached to a protein transduction sequence to promote cell entry is currently sold commercially as a TRAF6 inhibitor (e.g. Novus Biologicals NBP2-26506) [16]. The reported affinity of a RANK peptide with sequence RKIPTEDEY for TRAF6 is 78 µM, determined by isothermal titration calorimetry [19]. The same study reported a K_d_ of 84 µM for the CD40 TIM6 peptide, and we showed that peptides from our screen bind ∼10-fold tighter than the CD40 peptide (Figure S2). Thus, peptides from our screen, possibly further optimized by adding an optimal flanking sequence, can serve as higher-than-native-affinity inhibitors. Having a broad range of peptide sequences that can disrupt TRAF6 binding, as we have generated here, can support further efforts to develop inhibitors with desirable properties, such as low immunogenicity and cell permeability.

Similar trends to what we observed in this study of TRAF6 have been observed for PDZ domains. Phage-display selections for PDZ domain-binding peptides have been conducted using both random peptide libraries and a library comprised of peptides from disordered regions of the proteome [2,4]. Interestingly, similar to our TRAF6 library screen, the random-peptide library hits included more hydrophobic sequences than the hits from the native-sequence library, and these peptides often contained tryptophan. A PSSM based on the random library hits was used to identify candidate viral binders with better scores than human binders, and several viral peptides were found to bind with low-micromolar affinities to the PDZ domain of SCRIB, a protein that is targeted by human papillomavirus [55,56]. This work and our results for TRAF6 indicate that random peptide library screening can provide high-affinity ligands for human protein recognition domains that exceed the affinity of native ligands, offering a strategy to identify inhibitors to modulate signaling that mimic strategies used by viruses.

## METHODS

### Vectors, bacterial cells, and cloning

The expression constructs and cell surface display constructs are detailed in Table S1. The TRAF6 trimeric construct, here termed T6cc, consisted of residues 310-504 of human TRAF6 (including the MATH domain and coiled-coil trimerization domain), an N-terminal BAP tag for biotinylation, and a hexahistidine tag for purification. The construct was expressed using a pDW363 vector to ensure biotinylation. A monomeric TRAF6 construct lacking the trimerization domain and BAP tag, here termed T6m, consisted of residues 350–501 of human TRAF6 and a hexahistidine tag. The construct was expressed in a pDW363 vector, although the lack of a BAP tag ensured no biotinylation of this protein. SUMO-peptide fusion constructs contained a BAP tag, hexahistidine tag, and SUMO tag. The construct was expressed in a pDW363 vector to ensure biotinylation. *E. coli* strains BL21(DE3), DH5α, and MC1061 were used for protein expression, cloning, and surface display, respectively. For bacterial surface display of TRAF6-binding peptides, the eCPX vector designed by the Daugherty group [57] was modified at the C-terminus to append a FLAG sequence, a linker containing a SfiI site, and the CD40 peptide sequence, which included 9 residues resolved in the X-ray structure of CD40 bound to TRAF6 (PDB ID: 1LB6 [19]) plus 8 flanking residues on each side of this core region. The CD40-derived sequence used is PTNKAPHP<u>KQEPQEIDF</u>PDDLPGSNT.

### Mutant library construction

The library was constructed using primers (from IDT) with NNK codons included in positions marked ‘x’ the motif xxxPxExxx, such that the theoretical size of the library was 20^7^ = 1.28 * 10^9^ unique members. The variable sequence was flanked by SfiI restriction sites for cloning. In parallel, a linear vector for cell-surface display containing the constant sequence of the display construct, with SfiI sites matching the library insert, was amplified by PCR. The insert and linear vector fragments were purified by PCR Cleanup Kits (Genesee Scientific) before SfiI digestion. Following purification of the digested fragments, a 5:1 ratio of insert:vector was added to a 200 µL T4 DNA Ligase (New England Biolabs) reaction and then incubated for 16 hours at 4 °C. The ligated mixture was electroporated into fresh electrocompetent MC1061 cells in four separate transformations. Transformed cells were transferred into 10 mL warm Super Optimal Broth media with 20 mM glucose (SOC media) and incubated at 37 °C for 1 hour. The 10 mL culture was then added to 1L of LB + 25 µg/mL chloramphenicol and grown to an OD600 of 0.6-0.8 before centrifugation and resuspension in LB + 20 % glycerol for freezing for storage.

### Protein purification and preparation

T6cc was co-expressed in BL21(DE3) *E. coli* with the biotin ligase BirA (from the pDW363 vector) for 5 hours at 37 °C. The protein was then purified using Ni^2+^-NTA affinity chromatography followed by gel-filtration chromatography into a final buffer of 20 mM Tris pH 8.0, 150 mM NaCl, 5% glycerol, 1 mM DTT. Purified protein was concentrated before storing at -80 °C in aliquots for later use. Concentrations of T6cc are reported as monomer concentrations. For solution binding studies, a monomeric variant of TRAF6 (T6m) was expressed in Rosetta2(DE3) cells overnight at 18 °C and purified similarly to T6cc. T6m was purified into a final buffer of 50 mM Tris pH 8.0, 180 mM NaCl, 5% glycerol, and 1 mM DTT. SUMO-peptide fusion proteins were co-expressed in Rosetta2(DE3) *E. coli* with the biotin ligase BirA (from the pDW363 vector) for 5 hours at 37 °C. The protein was purified by Ni^2+^-NTA affinity chromatography followed by gel-filtration into a final buffer of 20 mM Tris pH 8.0, 150 mM NaCl, 1 mM DTT, and 10% glycerol.

### Magnetic bead presorting (MACS)

To generate TRAF6-bound beads, 2 mL of vortex-mixed Invitrogen DynaBeads™ Biotin Binder beads were incubated with T6cc (33 pmol biotinylated T6cc/10 µL beads) for 2 hours at 4 °C and then washed in PBS buffer, as described by Angelini et al. [58]. The TRAF6-decorated beads were then added to cultures of induced cells expressing the peptide library (induced with 0.2% w/v arabinose for 2 hours at 37 °C). After incubation for 3 hours at 4 °C, beads were magnetically isolated for 60 seconds before aspiration and replacement of PBS buffer. Beads were then gently shaken in the fresh buffer for 5 minutes at 4 °C. The bead wash cycle was repeated 7 times before beads were placed in LB media for growth overnight. 100 µl of the final growth stock was serially diluted on LB + agar + 25 µg/mL chloramphenicol plates. Colony-forming units were tabulated to back-calculate the number of cells in the MACS-sorted library, which yielded 1.42 * 10^5^ cells.

### Bacterial FACS preparation

For enrichment sorts and single-clone FACS cell surface titrations, 5 ml cell cultures were grown overnight at 37 °C in LB + 25 µg/ml chloramphenicol and 0.2% w/v glucose. The next day the culture cell density was measured by OD_600_, and approximately 3.25 * 10^5^ cells of each stock were isolated for new growth in 5 ml LB. Upon reaching an OD_600_ of 0.5 – 0.6, cells were induced with 0.2% w/v arabinose for 2 hours at 37 °C. Density was again measured, and cells were pelleted by centrifugation and resuspended in PBS + 0.5% BSA. Cells were then aliquoted into a 96-well Multi-Screen HTS® GV sterile filtration plate (2 × 10^7^ cells per sample) and washed with fresh PBS + 0.5% BSA. Cells were then incubated in 30 µL of αFLAG-APC [PerkinElmer] (prepared at a 100:1 dilution in PBS + 0.5% BSA) at 4 °C for 15 min. Next, cells were resuspended in 50 µL of TRAF6 solution (25 µL PBS + 0.5% BSA mixed with 25 µL of the chosen TRAF6 concentration) and incubated at 4 °C for 60 min. Following a wash with 200 µL PBS + 0.1 % BSA, cells were resuspended in 30 µL streptavidin-PE (SA-PE) [ThermoFisher] (prepared at a 100:1 dilution in PBS + 0.1% BSA) and incubated at 4 °C for 15 minutes. Cells were then washed in 200 µL PBS + 0.1% BSA, resuspended in another 200 µL PBS + 0.1% BSA, and placed on ice prior to FACS analysis or sorting. FACS analysis was performed using an HTS Canto II instrument and sorting took place on a FACS Aria III cell sorter (BD Biosciences). Sorted cells were collected in 1.5 mL microcentrifuge tubes containing 500 µL Luria-Bertani media with 25 µg/mL chloramphenicol.

### Single-clone titration experiments

For single-clone titration experiments samples for FACS analysis were prepared as described above using eight concentrations of TRAF6 for each clone: 0 nM, 3 nM, 10 nM, 30 nM, 100 nM, 300 nM, 1 µM, 3 µM. Binding curves were generated by plotting the mean PE value vs. TRAF6 concentration and fit to the following equation to determine a 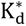 value:

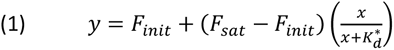

Where y is the mean PE fluorescence value and *x* is the concentration of TRAF6. F_init_, F_sat_, and K_d_ were treated as floating parameters; F_init_ is the *y* value in the absence of TRAF6 and F_sat_ is the *y* value at which the binding curve saturates. Although the cell-surface binding data fit well to a hyperbolic binding equation, this assay is not likely to be at equilibrium, and we discourage interpretation of the apparent binding constant 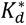 as a true equilibrium dissociation constant.

### Biolayer Interferometry (BLI)

BLI experiments were carried out on an Octet Red96 instrument (ForteBio). Streptavidin-coated tips (ForteBio) were pre-incubated for 10 min in BLI buffer (20mM Tris pH 8.0, 207 mM NaCl, 1 mM DTT, 1% Glycerol, 0.1% BSA, and 0.1% Tween-20). Biotinylated SUMO-peptides were immobilized on streptavidin tips. Loaded tips were then immersed in a solution of the TRAF6 MATH domain, which had been diluted to the relevant concentration in BLI buffer. Association data were collected at room temperature at an orbital shake speed of 1000 rpm (sampling rate) until the signal plateaued. Subsequently, TRAF6 bound tips were transferred to a well containing the above buffer, and dissociation data were collected until the signal plateaued. Due to the fast kinetics of the interaction, we elected to calculate K_d_ values using the steady-state signal of the association step. The raw association data of a SUMO-only control was subtracted from that of the SUMO-peptides. The normalized signal of the association step was averaged over 10 seconds after reaching a plateau and plotted against the concentration of TRAF6 MATH domain. The binding curve was fit to equation 1 in Kaleidagraph [59] using non-linear least-squares fitting to determine the dissociation constant.

### Enrichment sorting of MACS-presorted library

To isolate the best TRAF6 binders, we performed a five-round enrichment sort using the MACS-sorted library as the input. On each day, the library was sorted for TRAF6 binding as described above (*Bacterial FACS preparation*) using a single permissive gate set to collect successfully expressed TRAF6 binders. The gate was set manually each day using positive and negative binding controls. Selection for TRAF6 binding was gradually increased by using a lower concentration of T6cc each day (concentrations used: 300 nM, 100 nM, 30 nM, 10 nM, 3 nM). Collected cells were grown overnight before splitting half of the pool to continue the sort and the other half to harvest for plasmid DNA and subsequent Illumina sequencing. We performed two duplicate 5-day enrichment experiments, generating 10 total pools for deep sequencing

### Nonbinding clone FACS sorting

Using the unenriched (pre-MACS) library as input, a gate was drawn to define the region where peptide-expressing cells are found in the absence of TRAF6. This gate was used to collect 2 * 10^4^ cells in the presence of a high TRAF6 concentration ([T6cc] = 6 µM) to isolate clones with no detectable binding to TRAF6.

### Illumina amplicon preparation

Figure S3 gives an overview of this procedure. Sorted pools were grown overnight at 37 °C in LB, and bulk plasmid DNA was harvested by QIAprep miniprep kit (Qiagen). We then PCR amplified the variable region of the plasmids from each cell-sorted pool, appending a MmeI restriction site to the 5’ end. At the 3’ end, we appended: a) an unused, randomized 9 nt barcode UID sequence, b) a 6 nt indexing sequence for multiplexing (Illumina TruSeq), and c) a custom reverse-read annealing sequence. Barcodes are given in Table S1, amplicon construction is depicted in Figure S3, and a lookup table is provided in Table S2. Amplified fragments were a true digested with MmeI. A double-stranded DNA fragment with a 2 nt overhang matching the MmeI cut site was then ligated to each MmeI-cleaved fragment. This fragment contained the standard 5’ Illumina adapter sequence and one of 24 preselected 5 nt barcodes for sample multiplexing. 5’ and 3’ Illumina anchoring sequences were appended to the amplicons in a subsequent PCR amplification. More than 50 amplicons were Sanger sequenced (QuintaraBio) to assess amplicon quality, which revealed the expected sequences and variable positions. The sequencing length of each amplicon was 65 nt, so forward and reverse paired-end 40 nt reads overlapped by 15 nt. Immediately prior to Illumina sequencing, the MIT BioMicro Center verified fragment size for all pools by agarose gel and multiplexed all pools at equimolar amounts.

### Illumina data collection and processing

Illumina sequencing was performed on a NextSeq500. The reads were demultiplexed using custom python scripts: https://github.com/jacksonh1/NGS_demultiplexing. Reads that did not exactly match one of the barcode/index pairs (first 5 nts of the forward read and first 6 nts of the reverse read, respectively) were discarded. Additionally, we required each of the first 5 nts of the forward read to have a Phred score of 20 or greater. Next, the ‘reformat.sh’ tool from the BBTools suite (Version 38.94) was used to de-interleave the paired-end reads and filter for reads with an average Phred score greater than or equal to 20 (using the parameter: ‘minavgquality=20’) [60]. In our dataset, the forward reads covered the entire variable region of the displayed peptide. Therefore, reverse reads were discarded after de-interleaving, and only the higher-quality forward reads were used for further analysis. For each sample, we used custom Python scripts to count the abundance of each sequence in each sample at the DNA level, using an alignment-based counting strategy. Here, the forward reads were aligned to a counting template sequence covering the variable region of the display construct: *********CCT***GAA*********CCGG, where * represents variable nucleotide positions. Sequences that mismatched 3 or more times to constant positions of the template (non * positions) were removed. Sequence counts were then further collapsed to just the TRAF motif region: *********CCT***GAA*********. The result was a list of sequences and their associated read counts for each sample. NGS data have been deposited at the GEO with the accession number GSE149328. Processed data files are provided as supplementary material (see supplementary information for file descriptions). All sequence logos in this study were generated using the logomaker python library [61].

### Enrichment data analysis

For each replicate enrichment experiment, we first removed any sequences that didn’t have 50 or more reads on at least one of the 5 enrichment days or the input library (MACS sorted library). The remaining sequences were translated into amino acid sequences, and only those DNA sequences coding for peptides matching the xxxPxExxx motif were kept for further analysis. Amino acid sequences containing the characters “*” or “X” were then removed (corresponding to sequences containing stop codons or “N” bases). Read frequencies were calculated by dividing the read count of each sequence in each sample by the total number of reads in that sample. When a sequence had fewer than 20 reads, the frequency was set to 0 to minimize effects from low read counts. To determine a set of TRAF6 binding sequences, we first filtered for sequences with 20 or more reads on day 4 and/or day 5 in either replicate enrichment. The resulting list of binders was then further filtered to include only those sequences that enriched 2 or more times (defined as an increase in read frequency from one day to the next day) during either enrichment replicate, yielding a final list of 236 unique TRAF6-binding peptides.

### Nonbinder data analysis

The NGS data from the nonbinder FACS sample (*Nonbinding clone FACS sorting*) were analyzed to define sequences of peptides that don’t bind to TRAF6. Sequences were filtered to include only those DNA sequences coding for peptides matching the xxxPxExxx motif and having a read count of 20 or more. The final list of nonbinders contained 1200 unique peptides.

### Generation of PSSM for proteome scanning

To generate a position-specific scoring matrix (PSSM) from the enrichment and nonbinder data, we used pLogo, which uses log-odds-based scoring to generate a PSSM from a given set of foreground sequences and background sequences [32]. We used the 236 unique TRAF6-binding peptides determined from the enrichment experiment as the foreground and the 1200 unique nonbinding peptides from the nonbinder sample as the background.

### Scoring motif matches in the proteome

To generate a table of TRAF6 motif instances in the proteome, we used SLiMSearch4 [33] to find all matches to the consensus TRAF6 binding motif (regex: “…P.E..[FYWHDE]”) in the human proteome. We used the SLiMSearch “shared functional annotations” feature to allow filtering the hits to proteins that share Gene Ontology (GO) terms with TRAF6. The set of GO terms used to filter the hits can be restricted by the likelihood that a given term is shared by any two proteins in the proteome (“sig” or “p-value”). We used this feature to create filters of different cutoff values (sig <= 0.01, 0.001, 0.0001, and 0.00001) to allow filtering hits to those that share TRAF6 GO terms with sig less than or equal to the given cutoff value. The HIPPIE database was used to label motif instances in proteins that are annotated to interact with TRAF6 [35]. AlphaFold 2 structure predictions for the human proteome were downloaded from the AlphaFold Protein Structure Database [36–38].

### Selection of binding and nonbinding peptides for modeling studies

48 binders were chosen from the binding sequences identified in the enrichment experiment for structure-based modeling. 3 of the 48 binders (RNVPEESDF, RNVPEESTW, and WNMPAEYDF) came from an earlier analysis of the enrichment data and are not present in the final set of 236 binders. However, all 3 sequences enrich at least once during the enrichment experiment and are likely real binders despite not making the final cutoff. Additionally, 41 nonbinder sequences were selected from the nonbinder pool for structural modeling.

### Computational Rosetta modeling

The pipeline for modeling mutated peptide interactions proceeded as follows. First, all structures were alchemically mutated onto the crystal structure of TRAF6 bound to the CD40 peptide (seq: KQEPQEIDF, PDB: 1LB6) using FoldX. For each mutated pose, we used Rosetta relax to remove steric and angle violations. Next, the Rosetta FlexPepDock module was used to create 500 poses of each using the lowres_preopt flag to more aggressively sample the space in case of necessary residue rearrangement. The talaris2013 score function was used for all model scoring in Rosetta. The top pose by Rosetta score was isolated from each mutated sequence and used to rationalize residue preferences for both strong and weak binders. The Rosetta version used was rosetta_bin_linux_2017.08.59291_bundle.

### Scoring peptide binding affinity

We implemented two different computational pipelines for scoring peptide binding to TRAF6: FlexPepBind (FPB) [30] and an in-house protocol based on computing a *detaching Temperature* (DetachT) by using short molecular dynamics simulations at increasing temperatures.

Structures were prepared using TRAF6-CD40 complex structure 1LB6 as a model. All sequences were 9 amino acids long and shared the PxExxAr short linear TRAF6-interacting motif (TIM6). We assumed that all peptides bound in the canonical TRAF6 binding groove with a position similar to that of the CD40 peptide KQE<u>PQE</u>IDF [19].

Binding energies were computed using the FPB program implemented in Rosetta version 3.6 with scoring function ref2015 [62]. We generated models of peptide-protein complexes starting with structure 1LB6 (chains A, TRAF6 MATH domain, and B, CD40-native peptide), first relaxing the structure with the Rosetta Fast-Relaxation protocol to remove internal clashes and any angle violations in the receptor and the native peptide. We then introduced point mutations into the CD40 peptide, keeping the backbone atoms fixed and optimizing the sidechain conformations of mutated residues using the Fixed-Backbone Design package with Resfile flag [63]. Next, the Rosetta FPB module was used to sample 100 variations of the docking pose for each peptide, allowing both backbone and sidechain atoms to move, using the refinement flag, and applying harmonic constraints around the crystallographic distances between the peptide and TRAF6 to reduce the conformational sampling space. Specifically, we restrained backbone hydrogen bond distances between the peptide residue in position (+0) and TRAF6 residue G472 and between the peptide residue in position (+2) and TRAF6-G470 G472 (using the observed distances in structure 1LB6). Models were ranked based on total interface score, calculated as the sum over energy terms contributed by interface residues of both partners. Interface residues were defined as those with Cβ (Cα for Gly) within 8 Å of any atom in the TRAF6 protein. We used the lowest-interface score complex for our analysis. The following is the command-line flag array for modeling peptides using Rosetta: ($name indicates a wildcard inserted to match the peptide to be run)

$rosdir/FlexPepDocking.static.linuxgccrelease -s

$name\_Dock_0001.pdb -native $nativepdb –

lowres_preoptimize -pep_refine -nstruct 500 -use_input_sc –

ex1 -ex2 -out:file:silent $name\_Dock.silent –

out:file:silent_struct_type binary

For the Detach-T protocol, the crystal structure was initially minimized and equilibrated for 20 ns with CHARMM36a using ACEMD code. The resulting structure was then mutated using the VMD-Mutator tool to introduce changes into structure 1LB6 [64]. We ran MD simulations with the temperature increasing from 300 K up to 1000 K, using a temperature step of 10 K every 100 ps for a total time of 5 ns, restraining protein CA that were more than 15 Å from the binding pocket to avoid protein diffusion in the unit cell. For each peptide, we recorded the temperature at which the distance between the geometrical center of TRAF6 residue 471 and the center of mass of the peptide segment composed of residues p0(PRO)-p1(variable)-p2(GLU) increased to greater than 7 Å.

### Molecular Dynamics simulation of a subset of TRAF6-binders complex

For 89 complexes, we performed short molecular dynamics (MD) simulations to identify key interactions or disruptive elements that influence peptide binding. Simulations were performed in NAMD using the CHARMM36m force field [65,66]. Each of 89 TRAF6-peptide complexes was solvated with a 15 Å pad of TIP3P water (resulting in a final simulation box of ≈80,000 atoms). Simulations were performed at a constant pressure of 1 atm and temperature of 300 K, a non-bonded cut-off of 12 Å, rigid bonds between heavy atoms and hydrogen atoms [67], and particle-mesh Ewald (PME) long-range electrostatics [68]. All complexes were first subjected to 1,000 energy minimization steps. Relaxed models were then equilibrated for 50 ns using a time step of 2 fs with all Ca atoms restrained by a 10 kcal mol^−1^ Å^−2^ spring constant. Finally, 100 ns production runs were done using ACEMD [69], with non-bonded cut-off and PME parameters set as in the equilibration phase, and the time step increased to 4 fs. To prevent protein diffusion in the water box, a restraining spring constant (5 kcal mol−1 Å−2) was applied to all Cα atoms of the protein more than 15 Å from the peptide-binding pocket.

Structures from the production runs were analyzed to determine root mean square deviations (RMSD), root mean square fluctuations (RMSF), and the presence/absence of specific interactions (hydrogen bonds, salt-bridges) using a Donor(D)-to Acceptor(A) distance cutoff of 3.2 Å; hydrogen bonds were additionally required to have an A-D-H angle of < 30°. We also checked for structurally favorable aromatic sidechain arrangements. In particular, cation-pi interactions were defined using the distance between the indole/phenyl group centroid and the guanidium centroid or amino group for Arg/Lys, respectively, and the angle between the respective planes. The angle was defined between the normal vectors to the planes of the sidechain rings, the guanidium group, or the positively charged groups. To qualify as cation-pi interaction, the distance had to be below 5.5 A. If the sidechains had an angle between 45 and 135 degrees, the cation-pi interaction was defined as T-shaped, otherwise as stacked [70,71]. We applied a similar definition for pi-pi interaction, setting the distance threshold between the centroids of the two aromatic rings to 7 Å, and the angle range between 75° and 90° for T-shaped or < 15° for stacked (parallel displaced or vertical) [72–74]. MATH domain charge distributions for Figure 5 were computed using the VMD-APBS module [75,76].

## Supporting information

enrichment_rep1_readcounts.csv

enrichment_rep2_readcounts.csv

enrichment_rep1_readcounts-processed.csv

enrichment_rep2_readcounts-processed.csv

nonbinder_readcounts-processed.csv

final_binder_list.txt

MD_sequences.xlsx

## Author contributions

A.E.K. conceptualized the project with D.W., F.R., J.C.H. D.W. performed library screening, next-generation sequencing (NGS) prep, and FACS experiments. V.S. wrote code for raw NGS data processing. J.C.H. processed and analyzed the NGS and FACS titration data (Figures 2, 6, and S1; supplementary data files). F.R. performed the molecular dynamics simulations (Figures 3, 4, and 5). J. C. H. performed the proteome analysis (Figure 6 and Table S3). A S. performed the BLI experiments (Figure S2). J.C.H. and A.E.K. wrote the manuscript with input from all authors.

