## Supplementary information for "Molecular determinants of TRAF6 binding specificity suggest that native interaction partners are not optimized for affinity"

#### Extended methods

##### Stepwise protocol for amplicon generation of Illumina substrates

Sorted pools from the enrichment and nonbinder experiments were grown overnight in 10 mL LB + 25 µg/mL chloramphenicol (OD600 > 1.0) and then plasmid DNA from each pool was isolated (QIAprep miniprep kit using manufacturer's instructions). The resulting DNA pools were subjected to the following procedure (see Figure S3):

1. PCR1 amplified the variable region of the bulk plasmid DNA and attached a 5′ top strand overhang containing 12 nt of flanking DNA and a TCCACC MmeI recognition sequence. MmeI cuts 20 nt 3′ of its recognition site on the top strand and 18 nt 5′ on the bottom strand. The MmeI site appended in PCR1 in our construct was designed such that MmeI digestion results in a bottom strand 3′ overhang of 'AG' to match the overhang of our DNA adapters. If other adapter is chosen, make sure to generate the appropriate overhang. The top strand 3′ primer appends both a 9 nt unique identifier (UID) and a 6 nt index sequence. The appended UIDs in our analysis were not used in this work. After 5 cycles at $T_a$ = 60 °C, the next 20 cycles were run at $T_a$ = 66 °C such that only full-length fragments were reproduced. Phusion High-Fidelity Polymerase in HF Buffer was used (0.5 µL Phusion/50 µL reaction) for all PCR steps. The 6 nt index sequences that we used are indicated in Table S1. PCR products were purified with the Zymo DNA Clean and Concentrate Kit (Genesee).

| **PCR Recipe:** | **Thermal Cycler:** | |
|---|---|---|
| 10 µL Phusion HF Buffer 5X | 1. 3 min | 98 °C |
| 1 µL dNTPs (10 mM each dNTP) | 2. 30 sec | 98 °C |
| 1.1 µL PCR1a fwd primer (10 µM) | 3. 30 sec | 60/66 °C* |
| 1 µL PCR1a rev primer (10 µM) | 4. 30 sec | 72 °C |
| 10 µL Ligated DNA template | 5. Repeat steps 2–4 | 14X |
| 1 µL Phusion Polymerase | 6. 3 min | 72 °C |
| 25 µL water | 7. Hold | 4 °C |
| 50 µ total | | |

*: After 5 cycles with $T_a$ = 60 °C, add 1.5 µL PCR1b reverse primer (10 µM) and resume with $T_a$ = 66 °C

2. MmeI (NEB) was used as in the NEB instructions to cleave PCR1 products. The resulting DNA fragments should have a ~200 ng PCR1 product, and are used as input DNA for each digestion in the following recipe:

| **Digestion Recipe:** | **Thermal Cycler:** | |
|---|---|---|
| 9 µL PCR1 product | 1. 60 min | 37 °C |
| 2 µL CutSmart 10X buffer | 2. 20 min | 80 °C (heat deactivation) |
| 1 µL SAM | 3. Hold | 4 °C |
| 6 µL water | | |
| 2 µL MmeI (4U) | | |
| 20 µL total | | |

T4 DNA Ligase (NEB) was used to anneal adapters (Table S1) assigned by pool identity. The adapters, which contain both the 5’ Illumina forward sequencing primer sequence and a 5’ 5-nt barcode, each having a ‘TC’ top-strand 3‘ overhang for annealing to the designed ‘AG’ bottom-strand 3’ overhang left by MmeI cleavage of purified PCR1 products. Ligated products were then run on a 1% Agarose gel containing 10 µL GelGreen/100 mL total gel volume and bands ~117 nt were excised and purified with the Zymoclean Gel DNA Recovery Kit. Purified products were eluted in 20 µL water.

| **Ligation Recipe:** | **Thermal Cycler:** | |
|---|---|---|
| 15 µL MmeI-digested DNA | 1. 30 min | 25 °C |
| 2 µL T4 DNA Ligase Buffer 10X | 2. 10 min | 65 °C (heat deactivation) |
| 2 µL DNA Adapter (6 µM) | 3. Hold | 4 °C |
| 1 µL T4 DNA Ligase (400 U) | | |
| 20 µL total | | |

3. PCR2 amplifies the purified ligated amplicons to add the 5’ Illumina anchor sequences to each strand of the amplicon. PCR2 products were purified with the Zymo DNA Clean and Concentrator Kit, and eluted in 20 µL water. Selected samples were then sent for Sanger sequencing (QuintaraBio) to monitor amplicon quality and appropriate barcoding.

| **PCR Recipe:** | **Thermal Cycler:** | |
|---|---|---|
| 10 µL Phusion HF Buffer 5X | 1. 3 min | 98 °C |
| 1 µL dNTPs (10 mM each dNTP) | 2. 30 sec | 98 °C |
| 3.1.µL Fwd primer (10 µM) | 3. 30 sec | 66 °C |
| 3.2.µL Rev primer (10 µM) | 4. 30 sec | 72 °C |
| 10 µL Ligated DNA template | 5. Repeat steps 2–4 | 14X |
| 1 µL Phusion Polymerase | 6. 3 min | 72 °C |
| 25 µL water | 7. Hold | 4 °C |
| 50 µL total | | |

### Supplementary Figures

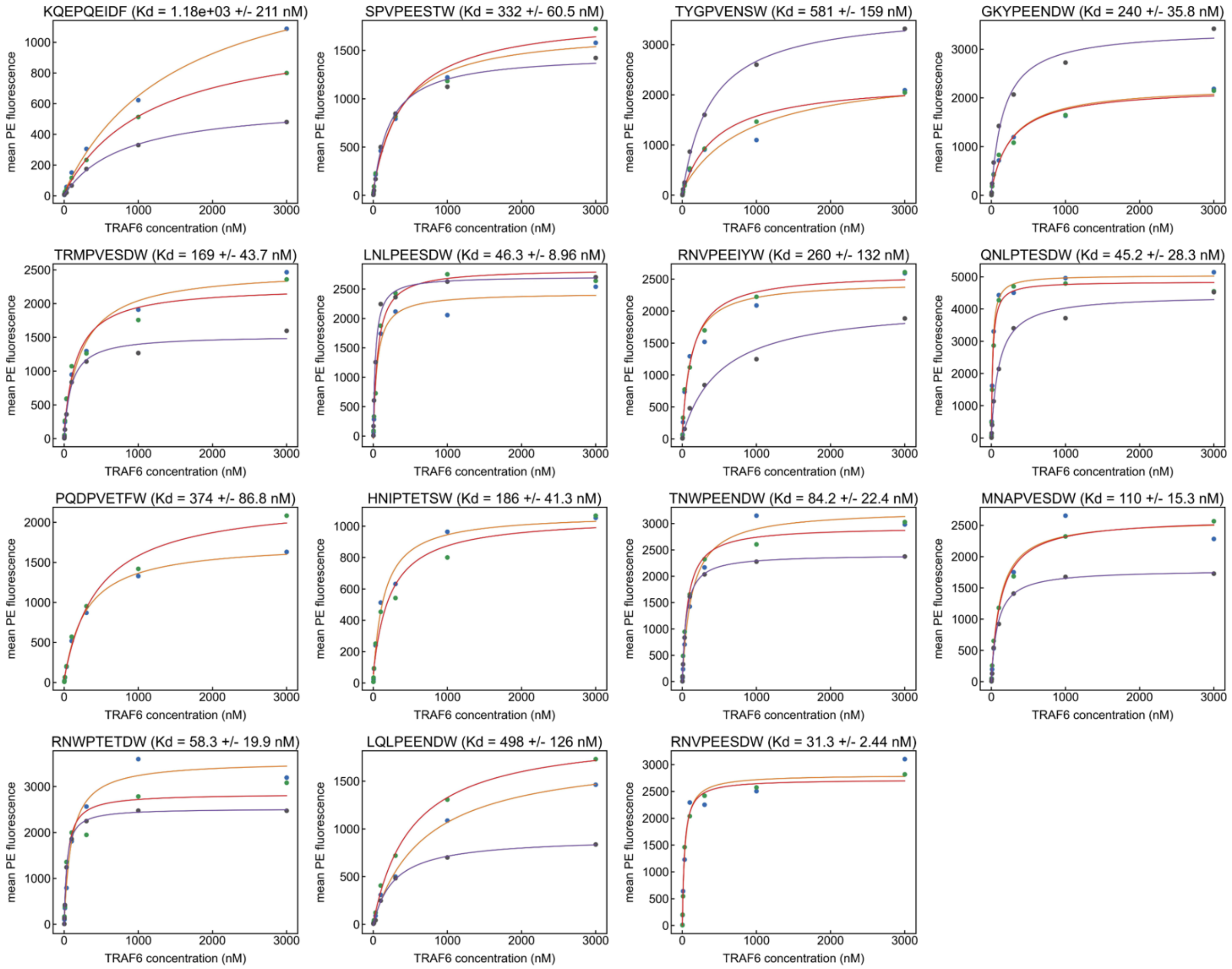


**Figure S1.** Binding curves from single-clone FACS titrations. Mean PE fluorescence is plotted against TRAF6 concentration and fit to a standard binding equation (equation 1). The mean $K_d^*$ from fitting each replicate independently and the associated standard error of the mean is shown above each graph.

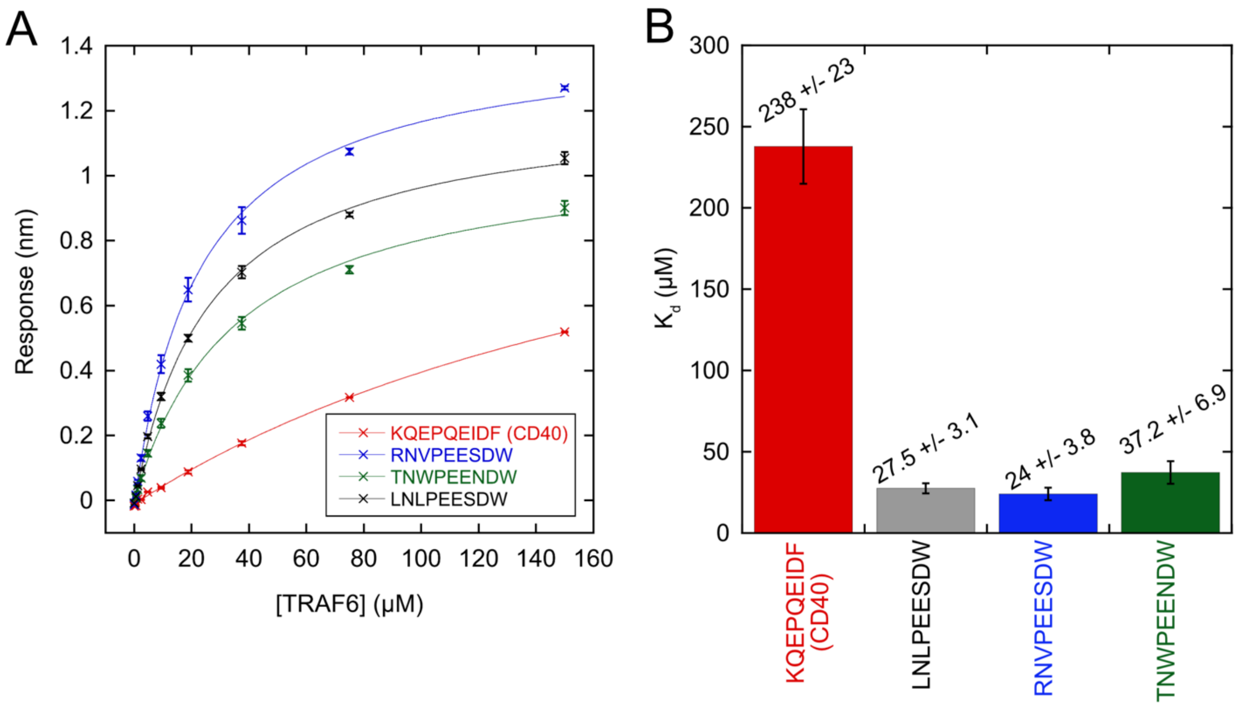


**Figure S2.** Biolayer interferometry (BLI) measurements of TRAF6 monomer (in solution) binding to different peptides (on tip). (A) Binding signal is plotted against TRAF6 concentration and fit to a standard binding equation (equation 1). Error bars are the standard error of the mean of 3 replicate measurements. (B) Average dissociation constant from 3 independently-fit replicate binding curves. The error bars are the standard error of the mean. Note that the dissociation constant for the CD40 peptide is an approximation as the highest TRAF6 concentration used was only 150 μM.

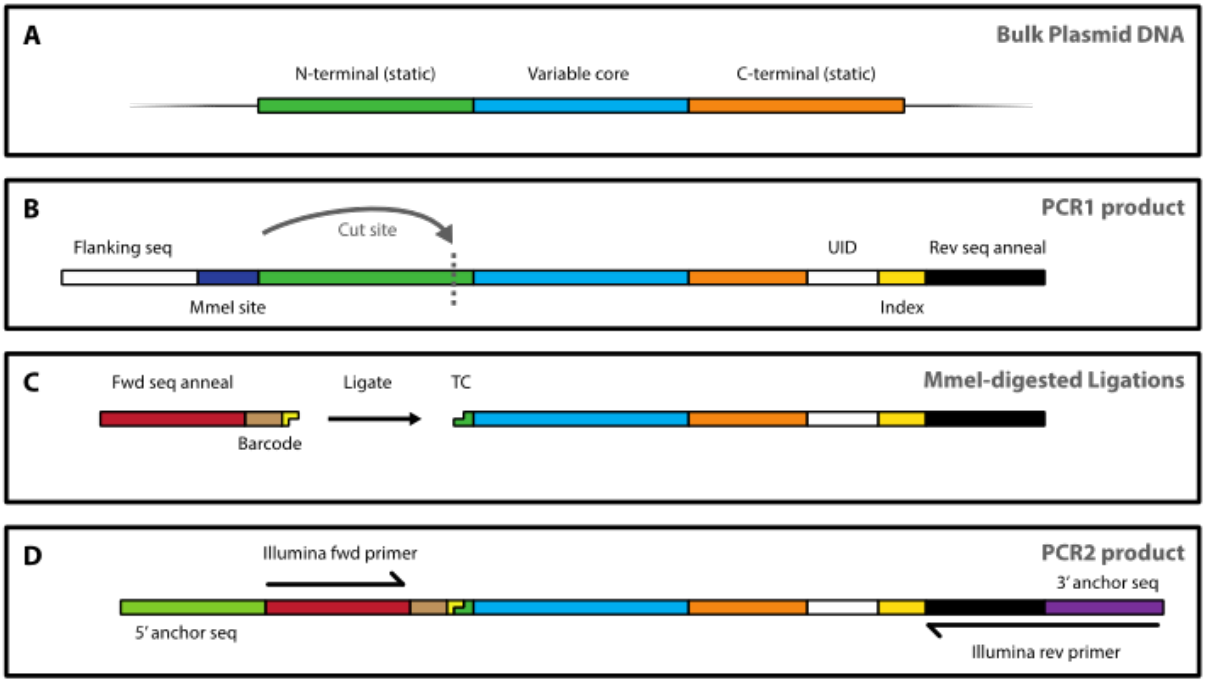


**Figure S3**. Schematic of amplicon preparation from sorted plasmid pools.

### List of provided supplementary data files

| Supplementary data file | Contents | column names | column description |
|---|---|---|---|
| enrichment_rep1_readcounts.csv | merged read counts for enrichment replicate 1 | pre-enrichment (MACSlib) | read counts in MACSlib (enrichment input) |
| | | day_1 | read counts after round 1 |
| | | day_2 | read counts after round 2 |
| | | day_3 | read counts after round 3 |
| | | day_4 | read counts after round 4 |
| | | day_5 | read counts after round 5 |
| | | seq | nucleotide sequence |
| enrichment_rep2_readcounts.csv | merged read counts for enrichment replicate 2 | pre-enrichment (MACSlib) | read counts in MACSlib (enrichment input) |
| | | day_1 | read counts after round 1 |
| | | day_2 | read counts after round 2 |
| | | day_3 | read counts after round 3 |
| | | day_4 | read counts after round 4 |
| | | day_5 | read counts after round 5 |
| | | seq | nucleotide sequence |
| MD_sequences.xlsx | sequences used in structural modeling studies | Binders | 48 binder peptide sequences |
| | | Nonbinders | 41 nonbinder peptide sequences |
| enrichment_rep1_readcounts-processed.csv | enrichment replicate 1 - read counts after filtering* | seq | nucleotide sequence |
| | | AA_seq | amino acid sequence |
| | | pre-enrichment (MACSlib) | read counts in MACSlib |
| | | day_1 | read counts after round 1 |
| | | day_2 | read counts after round 2 |
| | | day_3 | read counts after round 3 |
| | | day_4 | read counts after round 4 |
| | | day_5 | read counts after round 5 |
| enrichment_rep2_readcounts-processed.csv | enrichment replicate 2 - read counts after filtering* | seq | nucleotide sequence |
| | | AA_seq | amino acid sequence |
| | | pre-enrichment (MACSlib) | read counts in MACSlib |
| | | day_1 | read counts after round 1 |
| | | day_2 | read counts after round 2 |
| | | day_3 | read counts after round 3 |
| | | day_4 | read counts after round 4 |
| | | day_5 | read counts after round 5 |
| nonbinder_readcounts-processed.csv | nonbinder pool - read counts after filtering* final 1200 nonbinders | seq | nucleotide sequence |
| | | AA_seq | amino acid sequence |
| | | read counts | read counts |
| final_binder_list.txt | final set of 236 binder sequences | N/A | amino acid sequence |
| Supplementary_Table_S3.xlsx | TRAF6 motifs in the human proteome with PSSM score | columns explained at the top of the file | N/A |

*Filtering: Nucleotide sequences were collapsed to just the motif (*********CCT***GAA*********) and then translated into amino acid sequence. Peptides which did not match the TRAF6 motif (xxxPxExxx) or which contained a “*” or “X” character were removed. Any sequences that didn’t have a read count of 20 or more in at least one of the read count columns were removed.

#### Supplementary tables

**Table S1.** TRAF6 and peptide display constructs and index/barcode sequences used for multiplexing NGS samples.

| |
|---|
| **TRAF6 MATH-plus-coiled-coil construct - T6cc** |
| MA – (BAP tag) – GGSS – HHHHHH – GSGSGSM – (human TRAF6 residues 310-504) |
| ATGGCTGGAGGCCTGAACGATATTTTCGAAGCTCAGAAAATCGAATGGCACGAGGACACTGGTGGCTCGAGCCACCA<br>TCACCATCACCATGGTTCGGGCAGCGGATCGATGGATCATCAGATTCGTGAACTGACCGCGAAAATGGAAACCCAGAG<br>CATGTACGTGAGCGAACTGAAACGCACGATTCGTACACTGGAAGATAAAGTGGCGGAAATTGAAGCGCAGCAGTGCA<br>ATGGCATCTATATATGGAAAATTGGCAATTTTGGCATGCATCTGAAGTGCCAGGAAGAAGAAAAACCGGTGGTGATTC<br>ATAGCCCGGGATTCTATACGGGCAAACCGGGTTACAAACTGTGCATGCGTCTGCATCTTCAACTGCCGACCGCGCAGC<br>GTTGCGCCAATTACATCAGCCTGTTTGTGCATACCATGCAGGGCGAATATGATAGCCATCTGCCGTGGCCGTTCCAAGG<br>CACCATTCGTCTGACCATTTTAGATCAGAGCGAAGCGCCGGTGCGTCAGAATCATGAAGAAATTATGGATGCGAAACC<br>GGAACTGCTGGCTTTTCAACGTCCTACCATTCCGCGTAATCCGAAAGGCTTCGGCTATGTGACGTTTATGCACCTGGAA<br>GCTCTGCGCCAGCGTACCTTTATCAAAGATGATACCCTGCTTGTGCGGTGTGAAGTGAGCACCCGCTTTGATTAA |
| **TRAF6 monomer construct - T6m** |
| (human TRAF6 residues 350-501) – LE – HHHHHH |
| ATGAATGGCATCTATATATGGAAAATTGGCAATTTTGGCATGCATCTGAAGTGCCAGGAAGAAGAAAAACCGGTGGT<br>GATTCATAGCCCGGGATTCTATACGGGCAAACCGGGTTACAAACTGTGCATGCGTCTGCATCTTCAACTGCCGACCGC<br>GCAGCGTTGCGCCAATTACATCAGCCTGTTTGTGCATACCATGCAGGGCGAATATGATAGCCATCTGCCGTGGCCGTTC<br>CAAGGCACCATTCGTCTGACCATTTTAGATCAGAGCGAAGCGCCGGTGCGTCAGAATCATGAAGAAATTATGGATGCG<br>AAACCGGAACTGCTGGCTTTTCAACGTCCTACCATTCCGCGTAATCCGAAAGGCTTCGGCTATGTGACGTTTATGCACC<br>TGGAAGCTCTGCGCCAGCGTACCTTTATCAAAGATGATACCCTGCTTGTGCGGTGTGAAGTGAGCACCCTAGAACACC<br>ATCACCATCACCACTAA |
| **Peptides tested for cell-surface binding were displayed in an eCPX fusion construct with the following sequence:** |
| (eCPX residues 1 - 183 from ref. [1]) –<br>MKKIACLSALAAVLAFTAGTSVAGGQSGQSGDYNKNQYYGITAGPAYRINDWASIYGVVGVGYGKFQTTEYPTYKHDTSDY<br>GFSYGAGLQFNPMENVALDFSYEQSRIRSVDVGTWILSVGYRFGSKSRRATSTVTGGYAQSDAQGQMNKMGGFNLKYRY<br>EEDNSPLGVIGSFTYTEKSRTAS–GGGSGGGSDYKDDDDKGGGSGGGSGSGGQSGRGS–PTNKAPHP-xxxPxExxx-<br>PDDLPGSNT |
| ATGAAAAAAATTGCATGTCTTTCAGCACTGGCCGCAGTTCTGGCTTTCACCGCAGGTACTTCCGTAGCTGGAGGGCAGT<br>CTGGGCAGTCTGGTGACTACAACAAAAACCAGTACTACGGCATCACTGCTGGTCCGGCTTACCGCATTAACGACTGGG<br>CAAGCATCTACGGTGTAGTGGGTGTGGGTTATGGTAAATTCCAGACCACTGAATACCCGACCTACAAACACGACACCA<br>GCGACTACGGTTTCTCCTACGGTGCGGGTCTGCAGTTCAACCCGATGGAAAACGTTGCTCTGGACTTCTCTTACGAGCA<br>GAGCCGTATTCGTAGCGTTGACGTAGGCACCTGGATTTTGTCTGTTGGTTACCGCTTCGGGAGTAAATCGCGTCGCGC<br>GACTTCTACTGTAACTGGCGGTTACGCACAGAGCGACGCTCAGGGCCAAATGAACAAAATGGGCGGTTTCAACCTGAA<br>ATACCGCTATGAAGAAGACAACAGCCCGCTGGGTGTGATCGGTTCTTTCACTTACACCGAGAAAAGCCGTACTGCAAG<br>CGGAGGAGGCAGTGGTGGTGGGAGCGATTACAAAGATGACGATGACAAAGGCGGCGGTAGTGGGGGAGGCAGTGG<br>TAGCGGCGGCCAGTCTGGCCGTGGTTCTCCCACTAATAAGGCCCCGCATCCCAAACAAGAACCTCAGGAAATCGATTT<br>CCCGGACGATCTGCCGGGTAGCAACACATGATAA |
| **SUMO peptide construct used in BLI:** |
| MAG – (BAP tag) – DTGGSS – (TEV cleavage site) – HHHHHHH – GSGSG – (SUMO) – GSGSGGSGSG – xxxPxExxx |
| **Index and barcode sequences used to multiplex NGS samples** |
| Index sequences: ATCACG, CGATGT, TTAGGC, TGACCA, ACAGTG, GCCAAT, CAGATC, ACTTGA, GATCAG |
| Barcode sequences: ACTCG, ACTGT, AATGC, AGTCA, ATACG, ATAGC, CGATC, CTAAG, CTCGA, CGAAT, CTGGT, CGGTT, GACTT, GTTCA, GATAC, GAGCA, GATGA, GTCTG, TCGGA, TGACC, TACTG, TCCAG, TCGAC, TAGCT |

**Table S2**. fastq file sample key

| barcode file name | sample |
|---|---|
| barcode_0 | enrichment - replicate 1 - day 1 |
| barcode_1 | enrichment - replicate 1 - day 2 |
| barcode_2 | enrichment - replicate 1 - day 3 |
| barcode_3 | enrichment - replicate 1 - day 4 |
| barcode_4 | enrichment - replicate 1 - day 5 |
| barcode_6 | enrichment - replicate 2 - day 1 |
| barcode_7 | enrichment - replicate 2 - day 2 |
| barcode_8 | enrichment - replicate 2 - day 3 |
| barcode_9 | enrichment - replicate 2 - day 4 |
| barcode_10 | enrichment - replicate 2 - day 5 |
| barcode_20 | MACSLib pool* |
| barcode_5 | nonbinder pool |

*used as input for enrichment experiments
